# Binding profiles for 954 Drosophila and *C. elegans* transcription factors reveal tissue specific regulatory relationships

**DOI:** 10.1101/2024.01.18.576242

**Authors:** Michelle Kudron, Louis Gevirtzman, Alec Victorsen, Bridget C. Lear, Jiahao Gao, Jinrui Xu, Swapna Samanta, Emily Frink, Adri Tran-Pearson, Chau Huynh, Dionne Vafeados, Ann Hammonds, William Fisher, Martha Wall, Greg Wesseling, Vanessa Hernandez, Zhichun Lin, Mary Kasparian, Kevin White, Ravi Allada, Mark Gerstein, LaDeana Hillier, Susan E. Celniker, Valerie Reinke, Robert H. Waterston

## Abstract

A catalog of transcription factor (TF) binding sites in the genome is critical for deciphering regulatory relationships. Here we present the culmination of the modERN (model organism Encyclopedia of Regulatory Networks) consortium that systematically assayed TF binding events in vivo in two major model organisms, *Drosophila melanogaster* (fly) and *Caenorhabditis elegans* (worm). We describe key features of these datasets, comprising 604 TFs identifying 3.6M sites in the fly and 350 TFs identifying 0.9 M sites in the worm. Applying a machine learning model to these data identifies sets of TFs with a prominent role in promoting target gene expression in specific cell types. TF binding data are available through the ENCODE Data Coordinating Center and at https://epic.gs.washington.edu/modERNresource, which provides access to processed and summary data, as well as widgets to probe cell type-specific TF-target relationships. These data are a rich resource that should fuel investigations into TF function during development.

## INTRODUCTION

Precise deployment of gene expression across space and time drives the successful development and sustained health of all organisms. To understand the underlying causes of developmental disorders, polygenic diseases, and cancer, we must decipher the code that directs gene expression in diverse cell types and environments. Transcription factors (TFs) bind specific genomic sequences while interacting with regulatory proteins, RNA polymerase II, and the chromatin environment to direct gene expression programs. The set of TFs and their activity in a cell will determine which genes are expressed and to what level, thereby dictating cellular fate and function. Remarkably, despite the diverse body plans, lifespans and habitats of different species, the fundamental mechanisms by which TFs control gene regulation are highly conserved. Moreover, mutations in TFs are estimated to contribute to almost one-third of human developmental disorders (Vaquerizas et al. 2009), highlighting their central role in fundamental cellular processes.

A catalog of genomic sites bound by each TF (regulatory sequences) is therefore a critical step toward understanding gene regulation. Much of the work defining TF-chromatin interactions has been led by multi-lab consortia including ENCODE (Encyclopedia of DNA Elements) for humans and mouse (ENCODE Project Consortium et al. 2020) and modENCODE/modERN (model organism Encyclopedia of Regulatory Networks) for *C. elegans* and Drosophila (Gerstein et al. 2010; modENCODE Consortium et al. 2010; Boyle et al. 2014; Kudron et al. 2018; Celniker et al. 2009). These efforts have resulted in the systematic collection of millions of genome-scale measurements of TF binding, RNA transcripts, and chromatin accessibility and organization in a variety of organisms.

The nematode *C. elegans* and the fruitfly *Drosophila melanogaster* were selected for the modENCODE project in 2007 because of their experimental strengths and prominence as model organisms as well as for their compact, sequenced genomes, which simplifies identification of TF binding sites and candidate target genes. Critically, they also permit the mapping of TF binding in the living organism, which is not straightforward for human TFs. Many TFs in these two species are orthologous to human TFs. To a significant extent, they encompass the same broad classes of DNA binding domains, exhibit similar functions, respond to the same signaling inputs, and bind to similar consensus sequence motifs as human TFs. Over several decades, investigations of worm and fly TFs have illuminated their function during development and provided important insights into the function of human disease genes and human biology. Thus, studying key conserved factors in worm and fly will greatly enhance analysis, interpretation and the broader relevance of data gathered in ENCODE and other consortia-scale projects.

Working first as members of the worm and fly modENCODE consortia (modENCODE Consortium et al. 2010; Gerstein et al. 2010), and subsequently together as the modERN consortium (Kudron et al. 2018), we have generated hundreds of strains with tagged TFs, representing most of the TFs in each species. With these strains, we have characterized TF expression patterns and performed ChIP-seq analysis. These data sets define hundreds of thousands of binding sites throughout development for each species. Now, at the conclusion of this comprehensive effort, we report on the cumulative resources available to the community. We also analyze the global data to examine important TF features and relationships, including the correlation of co-binding by TFs at target genes, the frequency of high TF occupancy sites in the genome, and the impact of underlying sequence motifs and chromatin accessibility on TF binding. Finally, we begin to decipher tissue-specific codes of gene regulation for individual and groups of TFs using available single cell gene expression data in both species. Together the results demonstrate the value and richness of these available data, which will be crucial to further developing detailed network maps in model organisms. In turn, these maps will accelerate the understanding of how the cognate genes function in analogous networks in humans.

## RESULTS

### Strain Generation and Resource

The initial step in obtaining ChIP-seq data in our pipeline was to generate strains with GFP-tagged transcription factors. For this large scale project it was not feasible to generate TF-specific antibodies, and early experiments showed that representative GFP-tagged TFs rescued null mutants <u>(Gerstein et al. 2010; Venken et al. 2009)</u>.

To generate strains in the two species, we exploited the specific features and resources of each organism. For flies, available BAC clones that contained the TF and sufficient flanking sequence from two different libraries <u>(Venken et al. 2009)</u> were tagged using recombineering. Clones were introduced into flies using the ϕC31 integrase system and attP docking sites on either the second or third chromosome. PCR was used to confirm the location and presence of the targeted TF. As reported previously, these constructs generally can rescue their corresponding mutant phenotypes <u>(Venken et al. 2009)</u>. In the latter stages of the project, 151 lines were obtained commercially from Genetivision (https://www.genetivision.com/). Altogether strains were obtained for 616 TFs and 56 chromatin or nucleic acid binding proteins (**Table 1; Suppl. File 1**). The tagged TF strains capture all but 42 of the estimated 658 TFs (91.6%) in our curated list. The number of estimated TFs in the fly genome has fallen from 708 in 2013 (Hammonds et al. 2013), for a variety of reasons, including revised gene models and reassignment to other nucleic acid-associated proteins. Strains were not generated for TFs with complex genomic organization where there was no BAC containing the predicted TF transcriptional unit or when no MIMIC line (Venken et al. 2011) was available to produce a tagged TF.

**Table 1.**
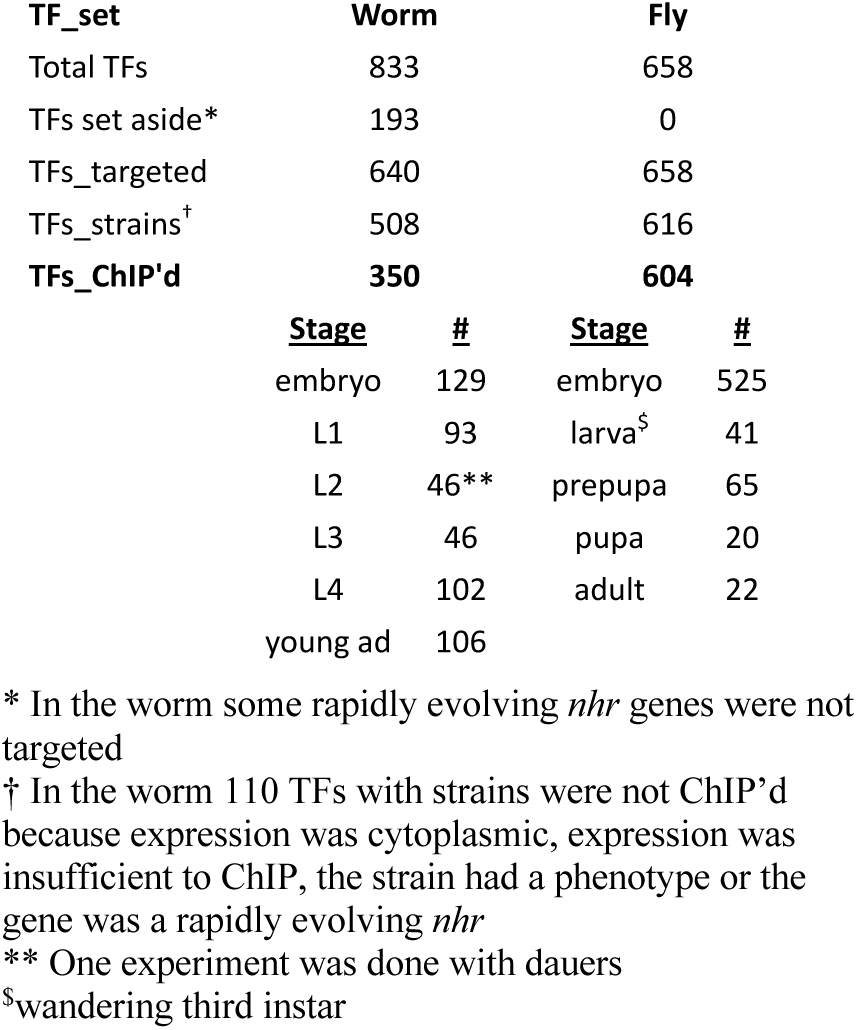

| TF_set | Worm | Fly |  |  |
| --- | --- | --- | --- | --- |
| Total TFs | 833 | 658 |  |  |
| TFs set aside* | 193 | 0 |  |  |
| TFs_targeted | 640 | 658 |  |  |
| TFs_strains<sup>†</sup> | 508 | 616 |  |  |
| **TFs_ChIP'd** | **350** | **604** |  |  |
|  | <u>Stage</u> | <u>#</u> | <u>Stage</u> | <u>#</u> |
|  | embryo | 129 | embryo | 525 |
|  | L1 | 93 | larva<sup>§</sup> | 41 |
|  | L2 | 46** | prepupa | 65 |
|  | L3 | 46 | pupa | 20 |
|  | L4 | 102 | adult | 22 |
|  | young ad | 106 |  |  |

\* In the worm some rapidly evolving *nhr* genes were not targeted
† In the worm 110 TFs with strains were not ChIP'd because expression was cytoplasmic, expression was insufficient to ChIP, the strain had a phenotype or the gene was a rapidly evolving *nhr*
\*\* One experiment was done with dauers
§wandering third instar

For worms, we primarily utilized the recombineered fosmid resource TransgeneOme to generate carboxy-terminal GFP tagged TFs <u>(Sarov et al. 2012)</u>. These tagged fosmids, generally selected so that the gene of interest was in the middle of the fosmid and flanked by other genes on both sides, were introduced into the worm by microparticle bombardment, which results in low copy (∼1-10) integration events at random sites in the genome. With only rare exceptions, the tagged strains have been viable and without readily detectable phenotypes. More recently, we switched to introducing the GFP directly into the endogenous gene via CRISPR for TFs that failed to yield strains by bombardment, that lacked a fosmid construct or that failed to produce a functional GFP-tagged TF upon recombineering <u>(Dokshin et al. 2018) (Ghanta and Mello 2020;</u> <u>Paix, Folkmann, and Seydoux 2017)</u>. The CRISPR approach has the advantage of being single copy and retaining the native regulatory context of the TF gene. To test whether this different mode of strain construction materially impacted TF function, we compared expression pattern and ChIP-seq results for fosmid-based and CRISPR-generated strains directly for five different TF genes and found no consistent differences (**Suppl. Table 1**). In total, strains with GFP tags were generated for 508 TFs and 45 nucleic acid-associated proteins. The total number of predicted worm TFs has fallen from an estimated 934 genes early in the project (Reece-Hoyes et al. 2005) to 833 total TFs in our curated list, due to factors such as revised gene models and reassignment to other nucleic acid associated proteins (**Suppl. File 1)**. Of the 833 total TFs, we targeted 642 and of these, 79% were successful, including 81 by CRISPR. Of the remaining 134 targeted TFs that lack strains, we have attempted to generate strains with either tagged fosmids or CRISPR without success.

These fly and worm strains are available through the Bloomington Drosophila Stock Center (BDSC) and the Caenorhabditis Genetics Center (CGC), respectively, and have been an extremely popular resource with the community. By 2023 the respective stock centers reported filling requests for 17,293 and 4,334 of these strains, respectively.

### ChIP-seq resource

The GFP tagged strains of both species were used to perform whole animal ChIP-seq using a highly specific, validated anti-GFP antibody (Kudron et al. 2018). In each case, synchronized populations of either flies or worms were grown to the desired stage, harvested and subjected to ChIP, using previously described methods (<u>(Gerstein et al. 2010)</u> <u>(modENCODE</u> <u>Consortium et al. 2010)</u>; see Methods). The bulk of fly experiments were performed on embryos because ChIP-seq data was more readily obtained at this stage and the majority of TFs were well expressed at this stage, while for the worm, the experiments were distributed more broadly across the life cycle and based on maximal expression predicted by RNA expression profiles and observed peak GFP fluorescence **(Table 1**). We elected not to perform ChIP-seq on some tagged worm strains because, based on prior experience, the TF was expressed in so few cells and/or at such low levels that a successful experiment was unlikely, or the expression was predominantly cytoplasmic. In a few cases in both worm and fly, despite repeated attempts, the data failed to meet quality criteria (see **Suppl. File 1** for tagged strains for which we did not obtain ChIP-seq data and the reasons). In total, we successfully performed ChIP on 604 fly TFs (92% of total TFs) in 674 different experiments and 350 worm TFs (55% of targeted TFS) in 519 different experiments (**Table 1**). In addition to the ChIP-seq data for TFs, we have also successfully generated ChIP-seq data for 45 additional DNA associated proteins in the worm in a total of 65 experiments, and 56 other DNA associated proteins in 64 experiments in fly. These results are all available at the ENCODE DCC and at https://epic.gs.washington.edu/modERNresource/, but the non-TFs are excluded from the further analysis presented here.

The resultant ChIP-seq data were processed through a peak-calling pipeline, using SPP (<u>(Kharchenko, Tolstorukov, and Park 2008)</u>; **Methods**). Data sets were immediately evaluated using the self-consistency ratio and rescue ratio parameters as defined by the ENCODE consortium (Landt et al. 2012). Passing data sets (see **Methods** for details) were submitted to the ENCODE Data Coordinating Center, until it stopped accepting submissions in October, 2022. Later data sets are only available at https://epic.gs.washington.edu/modERNresource/.

Because the SPP peak calling pipeline evolved over the life of the project, we reanalyzed all the data using a recent version of SPP and the DCC pipeline at the end of the ENCODE project to obtain more directly comparable datasets (see **Methods**). In these uniformly processed datasets, in the nuclear genome we detected 3,628,005 peaks in fly and 900,852 in worm, with a median of 4,416 peaks and 1,130 peaks per TF experiment respectively **(Figure 1 A, B)**. The basis for this considerable difference in peaks per experiment in the two species is unclear. The rightward distributions of peaks per experiment are broadly similar. Perhaps the fly, with no more TFs than the worm, but a much more complicated development, cellular composition and anatomy, achieves the added complexity by more extensive use of each TF. However, technical issues may play a role. For example, for fly we routinely collected three technical replicates, whereas for worm we collected only two. Reconstruction experiments suggest the use of the extra replicate could increase the number of peaks by about 20%. However, this amount is only a small fraction of the observed difference.

**Figure 1.**
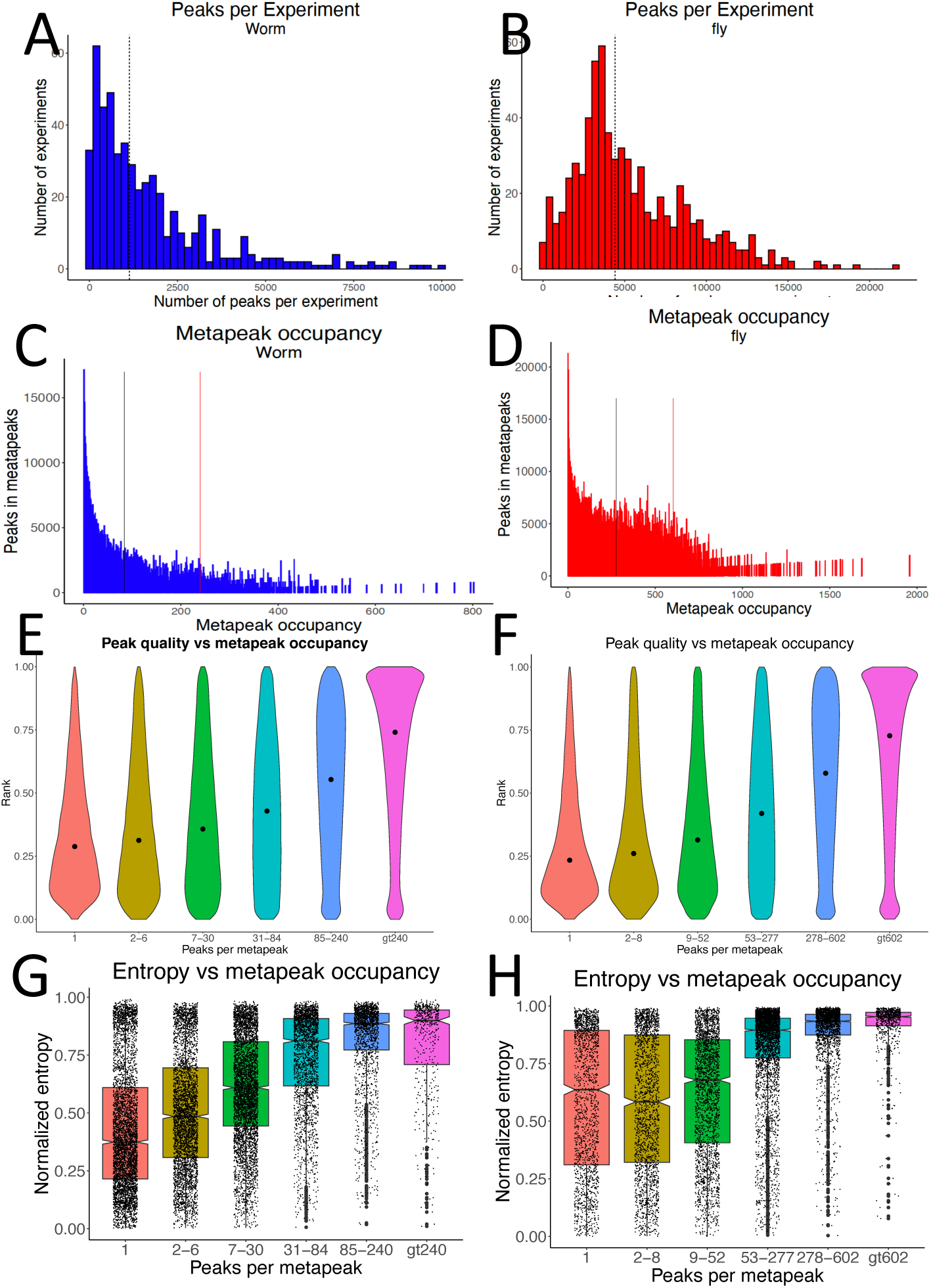
Peaks and Metapeaks. **A, B** Number of peaks per experiment (A) for the worm and (B) for the fly. Vertical lines denote the median number of peaks in worm (1,130) and fly (4,416). **C, D** The total number of peaks in metapeaks declines rapidly with increasing metapeak occupancy (the number of peaks in an individual metapeak) in both worm (C) and fly (D). The vertical bars indicate the thresholds used to define HOT (left) and ultraHOT (right) sites in each species (84 and 240 peaks respectively in the worm and 277 and 602 peaks in the fly. **E, F** In worm (E) and fly (F), the relative rank of the signal strength of peaks within metapeaks increases with metapeak occupancy. **G, H** Both worm (G) and fly (H) targets of high occupancy metapeaks show a predominance of high entropy genes (more uniform expression), while targets with lower occupancy metapeaks show lower entropy, indicating more cell type specific expression. The entropy of fly genes is shifted higher in the fly than the worm.

Most peaks in both worm and fly individually span much less than 1000 bases, with a median of 436 bases in fly and 376 bases in worm **(Suppl. Figure 1 A, B)**. Inspection of the few peaks more than 1,000 bases (4,518 in fly (0.12%) and 250 in worm (0.03%)), suggests that these often represent multiple overlapping peaks that SPP failed to separate.

### Peaks and metapeaks

As noted previously (Gerstein et al. 2010; Boyle et al. 2014), inspection of the mapped binding sites for each species revealed substantial clustering of peaks from multiple TFs in distinct regions around genes, with vastly differing numbers of peaks in each apparent cluster. To measure peak clustering in a principled manner, we treated each peak call as a sequence read and used MACS2 to call clusters of peaks, which for purposes here we have termed “metapeaks” (<u>(Zhang et al. 2008)</u>; see **Methods**). This process produced 77,969 metapeaks in the fly, including 19,136 singletons. For the worm there were 52,874 metapeaks, including 16,232 singletons. The span of these metapeaks increased only modestly with increasing numbers of peaks per metapeak (metapeak occupancy) with only 50 and 46 metapeaks spanning more than 3 kb in worm and 4 kb in fly respectively (**Suppl. Figure 1 C, D**). The DNA spanned by these sites amounted to 57,515,618 total bases in the fly (40% of the genome) and 27,564,528 bases in the worm (27% of the genome), even though the fly and worm have very similar number of bases in coding exons (33,650,573 for fly and 29,511,901 for worm). The greater coverage of the genome by the fly metapeaks could reflect the greater fraction of TFs assayed, the greater number of peaks per experiment, and/or greater complexity in the regulatory program. A single metapeak generally spans less than 3,000 bases in the fly and 2,000 bases in worm, even for metapeaks with high TF occupancy **(Suppl. Figure 1 C, D**), so that despite covering a large fraction of the genome in total, they represent quite discrete regions.

We investigated the overlap of the sites detected by ChIP-seq with recently obtained ATAC-seq data from both worm <u>(Jänes et al. 2018a)</u> and fly <u>(Cusanovich et al. 2018; Calderon</u> <u>et al. 2022)</u>and found good agreement (see **Methods**). We also compared the metapeaks with worm yeast-one-hybrid data <u>(Fuxman Bass et al. 2016)</u>, and, like the results of that paper, found less agreement between the different data sets, but the methods may well be assaying different aspects of gene regulation (see **Methods**).

The number of individual peaks (“occupancy”) within a metapeak declines rapidly, especially for the worm (**Figure 1 C, D**). Nonetheless, 1,408 metapeaks in fly contain more than 500 peaks and 491 metapeaks contain more than 250 peaks in the worm, suggesting regions of extremely dense TF binding. Highly Occupied Target (HOT) sites were recognized in early modENCODE papers <u>(Moorman et al. 2006; Gerstein et al. 2010)</u> and have been defined subsequently in various ways (Chen et al. 2014; Wreczycka et al. 2019). After exploring several approaches and comparing the HOT sites across the different stages (see **Methods**), we settled on the straight-forward criterion of defining the 5% most highly occupied metapeaks as HOT sites and the top 1% as ultra-HOT sites. For the worm, these thresholds are 84 and 240 peaks for HOT and ultra-HOT sites respectively and 277 and 602 peaks for fly. We found that most HOT/ultra-HOT sites were present across developmental stages in the worm, and even those that were not shared were usually only a few percent below the HOT/ultra-HOT threshold, indicating that most sites are quite consistent across development. Because the fly data are primarily from embryo, we did not examine HOT sites across development. For these reasons we combined the data from all stages to define HOT sites in each of the species for further analysis in this paper. Because of their high occupancy, these HOT/ultra-HOT metapeaks contain a disproportionate number of the individual peaks **(Table 2).**

**Table 2.**

| **Worm** |  |  |  | **Fly** |  |  |  |
| --- | --- | --- | --- | --- | --- | --- | --- |
| **Peaks per metapeak** | **Meta-peaks** | **Total peaks** | **Target genes** | **Peaks per metapeak** | **Meta-peaks** | **Total peaks** | **Target genes** |
| 1 | 17153 | 17153 | 2422 | 1 | 21332 | 21332 | 779 |
| 2-6 | 17913 | 58315 | 4291 | 2-8 | 25926 | 96908 | 2247 |
| 7-30 | 11369 | 162346 | 4868 | 9-52 | 16757 | 378090 | 2652 |
| 31-84 | 3823 | 193223 | 2465 | 53-277 | 10062 | 1264884 | 3953 |
| 85-240 | 2092 | 293067 | 1750 | 278-602 | 3109 | 1274688 | 2147 |
| > 240 | 524 | 176748 | 518 | > 602 | 783 | 592103 | 701 |

Because HOT sites may represent very open chromatin in many cell types that leads to non-specific TF binding, we sought to determine the relationship between metapeak occupancy and peak signal strength as reported by SPP. To normalize signal strength between experiments, we ranked the peaks within each experiment from 0 to 1, with 1 representing the strongest signal in any experiment. The distribution of peak ranks among metapeaks of differing TF occupancy shows the relative peak signal strength steadily increases with increasing cluster occupancy **(Figure 1 E, F)**. These results argue against HOT sites being locations of low-frequency, spurious binding and are consistent with the notion that they represent very open chromatin that facilitates strong TF binding (Partridge et al. 2020). The finding that singletons are enriched in the weakest binding sites may suggest that they are less likely to be functional.

We postulate that together these regions spanned by metapeaks sample much of the regulatory DNA in these two genomes. As such we would expect these regions on average to have conservation intermediate between coding exons and random unannotated regions of the genome. We initially used available multiple alignment groups for the two species, with the metapeaks regions of intermediate conservation. However, the differences in evolutionary distance between those species used in fly and those species used in worms (*C. elegans* has only quite distant relatives) made direct comparison of the results difficult. Instead, we opted to compare pairwise alignments between species selected to be of comparable evolutionary distance – *C. elegans* vs *C. remanei* (64% similarity) and *D. melanogaster* vs. *D. virilis* (62% similarity). In both species the regions of the genomes covered by metapeaks show levels of conservation intermediate between exons and random segments **(Figure 2A, B**).

**Figure 2.**
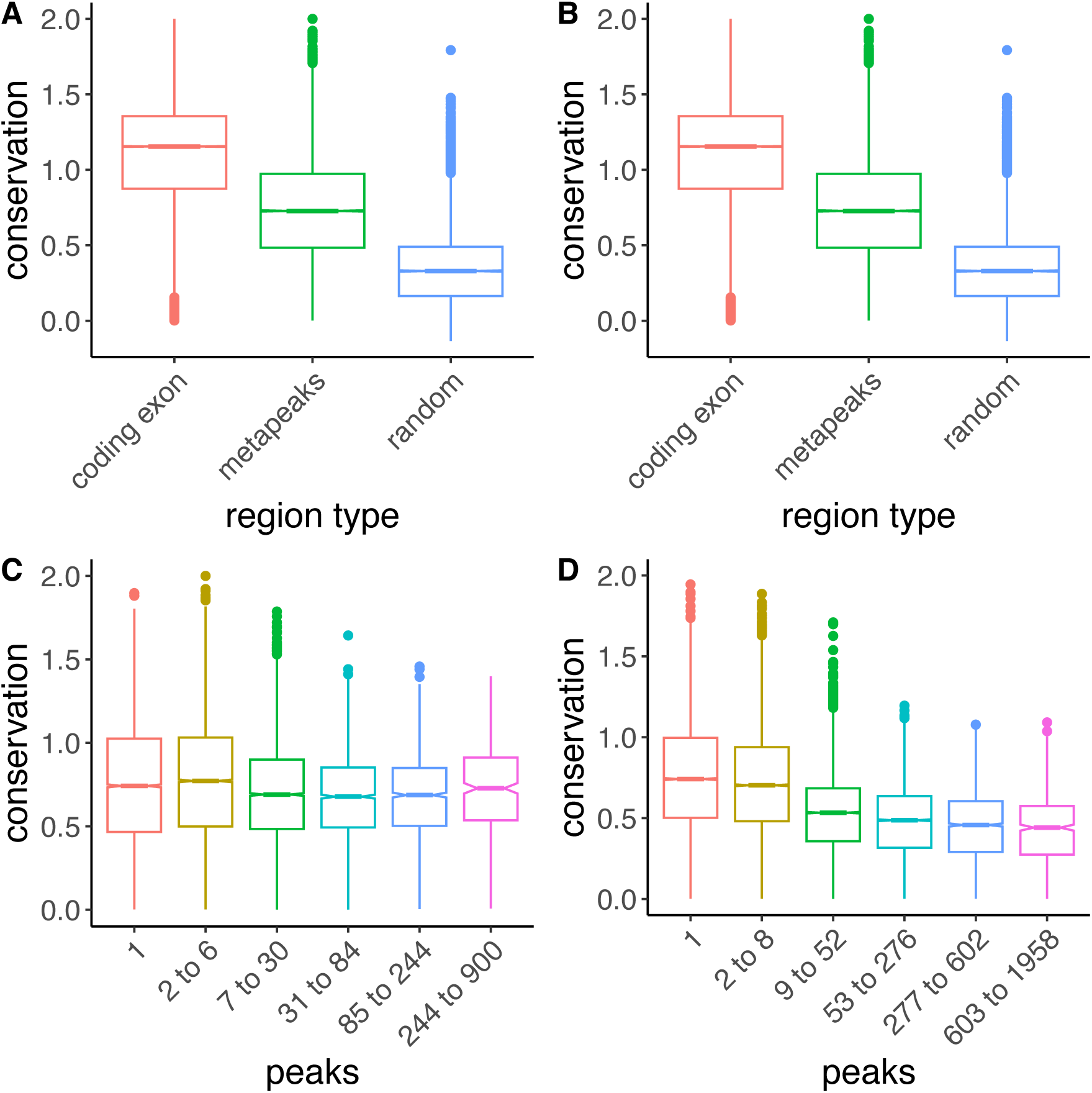
Conservation of metapeak regions. **A, B**. Regions of the genomes spanned by metapeaks show increased conservation compared to random regions but less conservation than coding exons. The fly exons are slightly less conserved than exon sequences in the worm. **C, D.** Metapeaks with increasing numbers of peaks show decreasing conservation, particularly in the fly.

We also evaluated the conservation of the DNA covered by the metapeaks based on the number of TFs per metapeak. In pairwise alignments, both species showed a decrease in conservation with increasing metapeak occupancy, with the trend more pronounced in flies **(Figure 2 C, D)**. Perhaps the larger span of the HOT and ultra-HOT metapeaks includes more unconstrained bases, with sequence-specific binding of only a subset of TFs critical to formation of the site, and the presence of many other less consequential TFs associated with open chromatin.

### Co-association of peaks in metapeaks

Inspection of the metapeaks suggested that certain TFs frequently occur together in the same metapeak. We therefore derived the Pearson correlation of each TF pair across the full set of metapeaks with TF membership greater than two, and clustered the results. When looking at all experiments together in the worm, TFs from the same stage often clustered together, as well as experiments with the same TF done on different stages. To avoid these stage-specific or TF-dependent effects we separated the experiments into three different groups for the worm (all embryo experiments, grouped larval stages L1-L3, and L4/young adult **(Figure 3, Suppl. Figure 2).** In each set, there are well-correlated TFs, including some known to be involved in muscle development (*hlh-1, rnt-1, unc-120*), and hypodermal development (*elt-1, elt-3, blmp-1*) in the embryo (**Suppl. Figure 2 A, B**) as well as other less familiar associations, for example in L4 and YA TFs likely involved in the gonad (*D2030.7, F33H1.4, ceh-91, madf-8, efl-1, madf-6*) and some likely involved in the intestine (*pqm-1*, *dve-1* and others)(**Figure 3A**).

**Figure 3.**
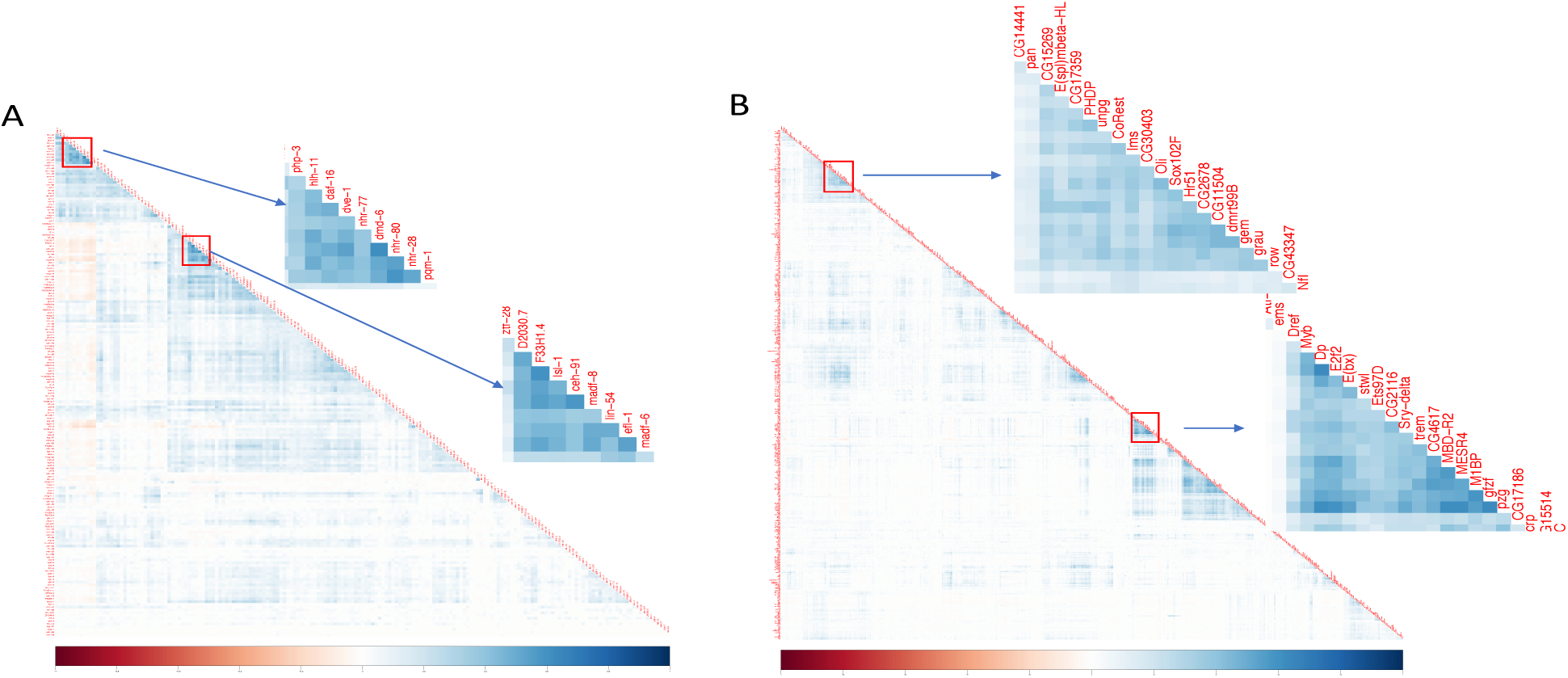
Correlation of TF-TF pairs. Peaks of some TF pairs occur in the same metapeaks more frequently than others as measured by Pearson correlation for both worm L4/young adult experiments (left) and fly experiments (right). Two clusters of TFs with correlated peaks are highlighted for each species but others are also evident. Negative associations are evident on the left side of the worm plot.

For the fly, we examined the Pearson correlation of the TFs in each of the stages assayed (**Figure 3B, Suppl. Figure 2 C, D, E**), but the relatively few experiments in the larva, pupa and adult stages limits their utility. However, the fly embryo has several high-density clusters, two of which are highlighted (**Figure 3 B**). The first cluster is composed of 21 genes of which there is molecular evidence of expression for 18 of the 21 (85%) in the nervous system, four only becoming active in larval and adult stages <u>(Hammonds et al. 2013; Calderon et al. 2022; Brown</u> <u>et al. 2014; Hongjie Li et al. 2022)</u>. The second cluster is composed of 23 TFs of which all are maternally deposited and 11 are maximally expressed in ovaries. These genes are members of chromatin insulator complexes (7 TFs; *Dref, E(bx), Sry-delta, M1BP, gfzf, pzg* and *ZIPIC*), embryonic development (5 TFs; *D, ems, Ets97D, MESR4* and *crp*), the dREAM complexes (4 TFs; *Dp, E2f2, Myb* and *CG17186*), chromatin associated proteins (3 TFs; *Atf-2, stwl* and *trem*) and one member of the NSL complex (*MBD-R2*). The dREAM and CP190 chromatin insulator complexes have been proposed to be required for transcriptional integrity, and it has been suggested that the dREAM complex functions in concert with CP190 to establish boundaries between repressed/activated genes <u>(Korenjak et al. 2014)</u>. The NSL complex regulates housekeeping genes <u>(Lam et al. 2012)</u>. Together the genes of the second cluster appear to be required maternally to set up general zygotic transcription.

### Motifs

To investigate the relationship between TF binding motifs and the peaks or metapeaks, we collected the motifs determined by *in vitro* experiments from the CIS-BP database (Weirauch et al. 2014). In both fly and worm, the number of *in vitro* motifs in a metapeak region is highly correlated with the number of peaks in the region (Figure 4 A,B), supporting the idea that HOT/ultra-HOT sites are not mere spurious binding. On average, *in vitro* motifs appeared in 30.6% and 12.9% of all the metapeaks in fly and worm respectively, without showing preference for singleton or HOT/ultra-HOT sites. We then inferred TF binding motifs for each TF from its ChIP-seq peaks and considered it as a successful inference if we observed any significant similarity between the best three inferred motifs and the corresponding *in vitro* motifs. Compared to using all the peaks, we observed an increase in the number of successes when we only used a subset of peaks after removing HOT and ultra-HOT sites and singletons (**Figure 4 C, D**). Interestingly, with a more stringent HOT site threshold, further improvement was achieved. Nevertheless, both subsets resulted in a higher success rate than using all or a fixed proportion of peaks, consistent across worm and fly.

**Figure 4.**
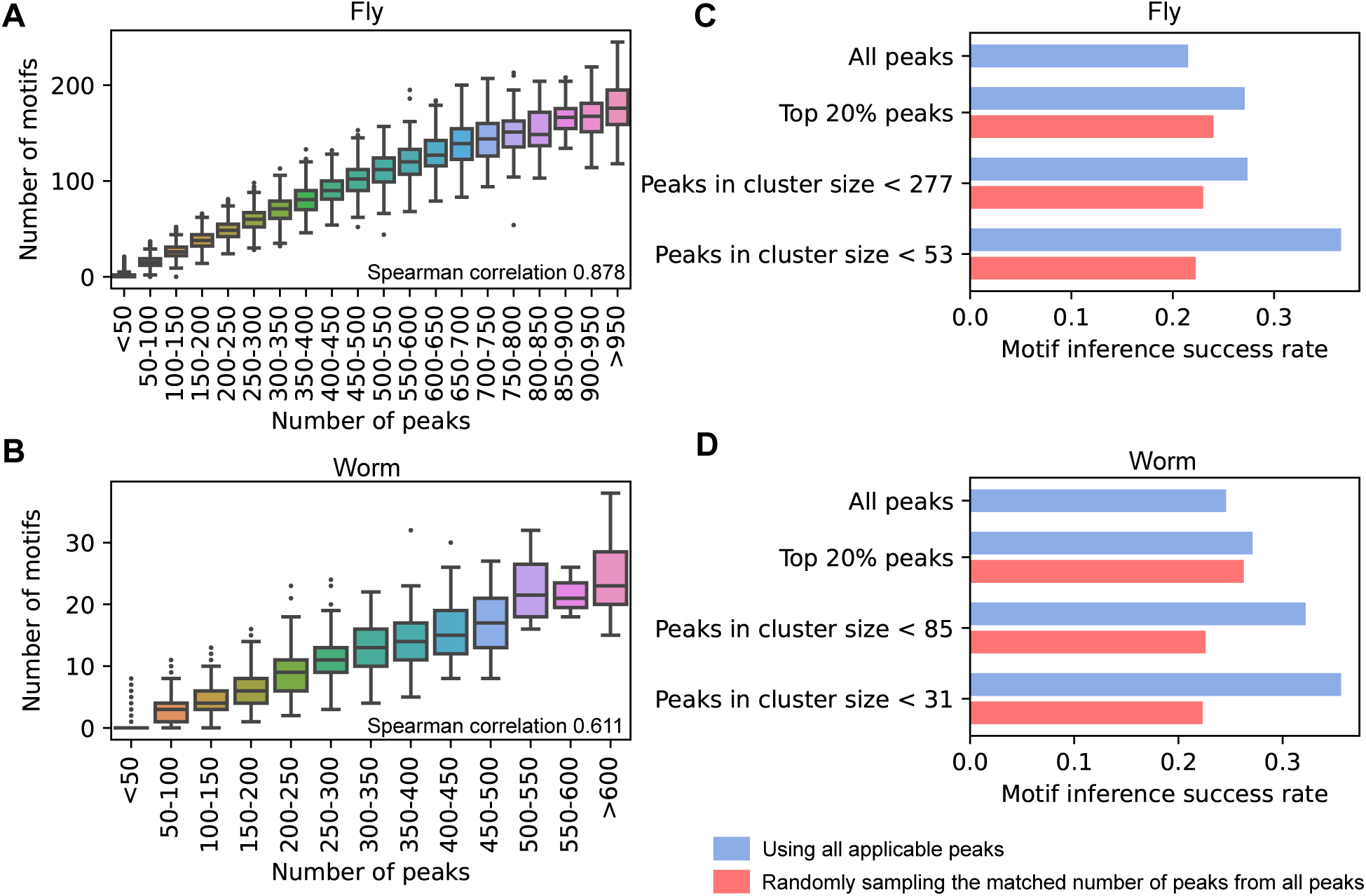
Motif analysis in the metapeaks. **A, B** The number of peaks in a metapeak region was highly correlated with the motifs in the region for both fly (A) and worm (B). The Spearman correlations for each are shown. **C, D** Utilizing the ChIP-seq data to obtain motifs, we found using a more stringent peak cluster threshold of 53 for fly (C) and 31 for worm (D) resulted in a better motif inference success rate compared to using all peaks, the top 20% of peaks or a larger cluster size. For each subset, we randomly sampled the same number of peaks from all peaks for better comparison (shown in red). This sampling process was repeated three times and the average value was shown.

This result suggested that excluding HOT/ultra-HOT sites may aid downstream motif inference. To test this idea, we inferred motifs for TFs that did not have *in vitro* motifs, using all peaks and only peaks left after stringent filtering of HOT sites filtering. We found that the inferred motifs were consistently found between these two inputs in 72.63% and 59.68% of experiments for fly and worm respectively. These inferred motifs could serve as important resources for further regulatory analysis.

### Distribution of peaks and metapeaks relative to protein coding genes

In order to evaluate the correlation between TF binding sites and mRNA abundance as measured in other experiments, we assigned the binding sites to candidate target genes. Binding sites can be at considerable distance from the genes they regulate ((Levo et al. 2022) and references cited in this paper), but most genes do have promoter proximal binding sites, and in these compact fly and worm genomes, transgenic reporter experiments indicate endogenous expression patterns can usually be produced from promoter proximal DNA (≤1-2kb). Exploiting this simplification, we assigned the apices of peaks and metapeaks to their nearest TSS as the primary target gene. In most cases we also assigned the next closest gene as an alternate target (see **Methods**). Similarly, we assigned metapeaks to target and alternate target genes.

A total of 13,109 genes (93.7% of 13,986 total) in fly and 16,822 genes (84.2% of 19981 total) in worm had at least one associated peak, with the number of associated peaks varying widely (**Suppl. Figure 1 E, F**). This variability is also reflected in the occupancy of the metapeaks associated with target genes. For worms 2,268 genes had one or more associated HOT or ultra-HOT sites and 2,422 genes had only singletons, with 1,715 having only one associated singleton (**Table 2**), while fly had 2,848 genes with associated HOT/ultra-HOT sites and only 779 with only singletons, with 494 having only one singleton.

The position of the peaks relative to their targets also varied. Both worm and fly exhibited a strong bias for peaks within 500 bases of the nearest TSS, with substantially more sites upstream than downstream of the TSS. The peak of the distribution in *C. elegans* is slightly farther upstream of the TSS than in Drosophila, perhaps reflecting the fact that for many *C. elegans* gene models the annotated TSS is actually the site of trans-splicing, with the start of transcription occurring further upstream (Allen et al. 2011; Saito et al. 2013). **(Figure 5 A, B)**. However, the distribution of peaks relative to the TSS varies for individual TFs, with some having a median position as much as 1 kb upstream, and others several hundred bases downstream of the TSS.

**Figure 5.**
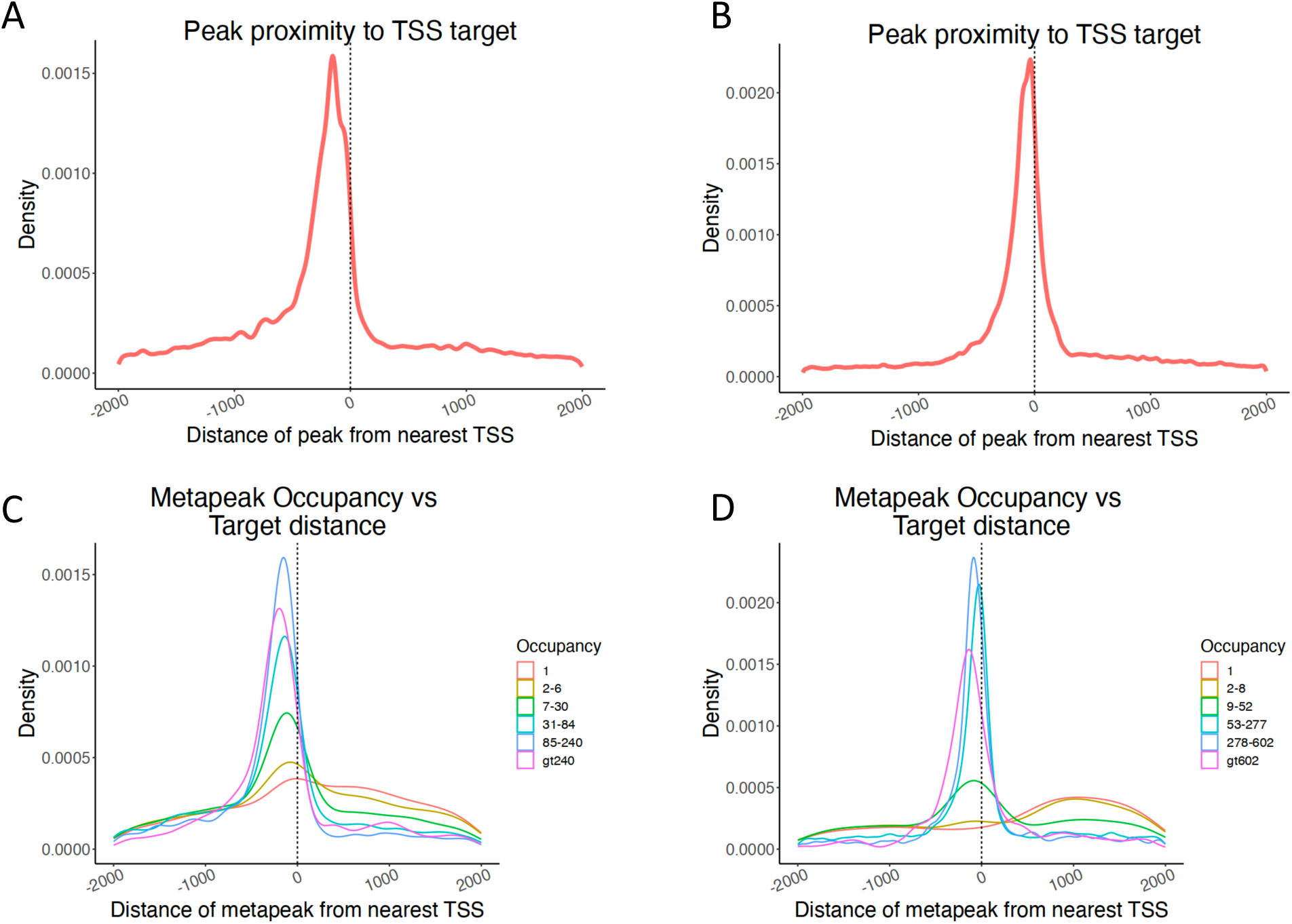
Peak position relative to TSS. **A, B** Peaks in worms (A) and flies (B) predominantly lie close to the TSS of the nearest gene. Worm peaks are slightly farther from the TSS, possibly reflecting the use of splice leaders in worm transcripts, so that the actual start of transcription is further upstream. The hint of two different distributions may reflect those genes with a splice leader and those without. **C, D.** Metapeaks with few peaks – less than or equal to six in the worm (C) and less than or equal to eight in the fly (D) – are more broadly distributed.

We next examined the relationship of the distance of metapeaks to TSS’s with respect to the number of peaks within the metapeak. High occupancy metapeaks were more tightly distributed near the TSS, with singletons and low occupancy metapeaks much more broadly distributed **(Figure 5 C, D)**. In fact, in flies, singletons and the lowest occupancy metapeaks were more prevalent downstream of the TSS. If position relative to the TSS is an important indicator of function, this distribution might suggest that singletons and very low occupancy metapeaks are less likely to be functional, but it could also reflect that active promoters tend to be so occupied with metapeaks that there is little real estate available for singletons.

### Complexity of regulation

Many genes in both worms and flies h ave complex expression patterns, with expression in multiple cell types at distinct times in the life cycle. This complexity is achieved by particular combinations of TFs in the different cell types at different times. To examine the relationship between TF binding and target gene expression in the worm, we exploited available single cell expression profiles (Packer et al. 2019; Cao et al. 2017; Calderon et al. 2022). For the fly data set, in order to increase the number of cell types, we removed the very young cells and remapped the remaining and reannotated the resulting clusters, obtaining 83 different cell types (see **Methods** for details). In order to estimate complexity of expression, we calculated the entropy of each target gene’s expression across cell types and time bins, where entropy here is defined as the sum of the negative fractional expression of a gene multiplied by the log_2_ of the fractional expression across all cell types. The entropy of a gene with absolutely uniform expression is log_2_ of the number of cell types, and the entropy of a gene with expression limited to just one cell type will be 0 (see **Methods**). We then subdivided the genes into those with ultra-HOT sites, those with HOT sites, etc., down to singletons and assessed the entropy of target genes. Genes with ultra-HOT or HOT sites had the highest entropy, reflecting broad expression, while genes with only low occupancy sites had low entropy, with expression in only a limited number of cell types or time bins (**Figure 1 G, H**). Broad expression might then be the result of the sum of many TFs acting on a target gene. An alternative explanation for such enrichment of high entropy genes with high-occupancy sites might be that broadly expressed genes have a more open chromatin configuration, allowing the binding of many factors, even if they don’t play a role in regulation of that target. Likely, both explanations contribute. By contrast, target genes with more cell type specific expression are associated with smaller metapeaks.

We also examined the relationship between the number of peaks associated with a gene and the number of clusters. Some targets have hundreds of associated peaks distributed in only one or two metapeaks, while others have multiple peaks distributed over many metapeaks (**Suppl. Figure 3 A, B**). As might be expected, housekeeping genes (Ghaddar et al. 2023) have predominantly large numbers of peaks in relatively few metapeaks (**Suppl. Figure 3 C**). These results echo the recent findings in yeast (Rossi et al. 2021). In contrast, genes with multiple transcription start sites (TSSs) tend to have a large number of associated metapeaks of variable occupancy (**Suppl. Figure 3 D, E**). The TF genes themselves are widely scattered in these plots, with some having only few associated peaks in one or two metapeaks, while others have multiple TSSs and multiple associated metapeaks, e.g., *ceh-38*, *daf-12*, *daf-16*, *egl-44*, *syd-9*, and *unc-62* in worms. The regulation of genes with multiple highly occupied metapeaks is likely to be very complex, with different regulatory sites driving expression in different cell types and times.

### Target expression patterns can reflect TF expression patterns

A central question for the ChIP data is to what extent, if any, the detected peaks or binding events, for each TF drive gene expression of its targets. Addressing this question is complicated by the observation that TF ChIP data often detect binding at many more sites than expected based on known or experimentally validated functional sites (Cusanovich et al. 2014; Hu, Killion, and Iyer 2007). Nonetheless, for strong, positive regulators the expectation would be that the expression patterns of TFs would correlate with the expression of targets, whereas negative regulators would be negatively correlated. The emergence of single cell RNA-seq data (scRNAseq) for both worm and fly offers an opportunity to investigate these relationships directly. For the worm we used scRNAseq data from the embryo, both terminal cells and their progenitors (Packer et al. 2019), L2 (Cao et al. 2017; Packer et al. 2019) and young adult data (Ghaddar et al. 2023). For the fly, since most of the ChIP data was collected in embryos we focused on the recent embryo scRNAseq data set (Calderon et al. 2022). Because the resolution of cell types in the published fly data set was limited, we subclustered the older cells, remapped them and assigned additional and more specific cell types, detecting 83 cell types in all (see **Methods**).

As a direct test of the relationship of the TF and its target expression patterns we aggregated the expression values for each target, summing the values across all cell types. We matched the ChIP data with single-cell expression data of related stages. For TF-target pairs in the worm we used ChIP data from the embryo with the embryo expression data, ChIP data from L1, L2 and L3 with the L2 expression data, and ChIP data L4 and young adult with the young adult data. For the fly, we only matched the embryo ChIP data with the embryo expression data, since most of the ChIP data was done in embryos.

We used a mean-centered expression matrix to evaluate TF-target expression. As examples, for *blmp-1* in worms (**Figure 6 A, B**) or GATAe in flies (**Figure 6 C, D**), this results in negative values for cell types with no or very low expression, but still high positive values for other cell types. For genes with broad, uniform expression, most values will be around zero, thus minimizing their impact on the aggregate pattern. For targets, we only considered targets that were not associated with HOT sites, recognizing that these genes with HOT sites were generally broadly expressed.

**Figure 6.**
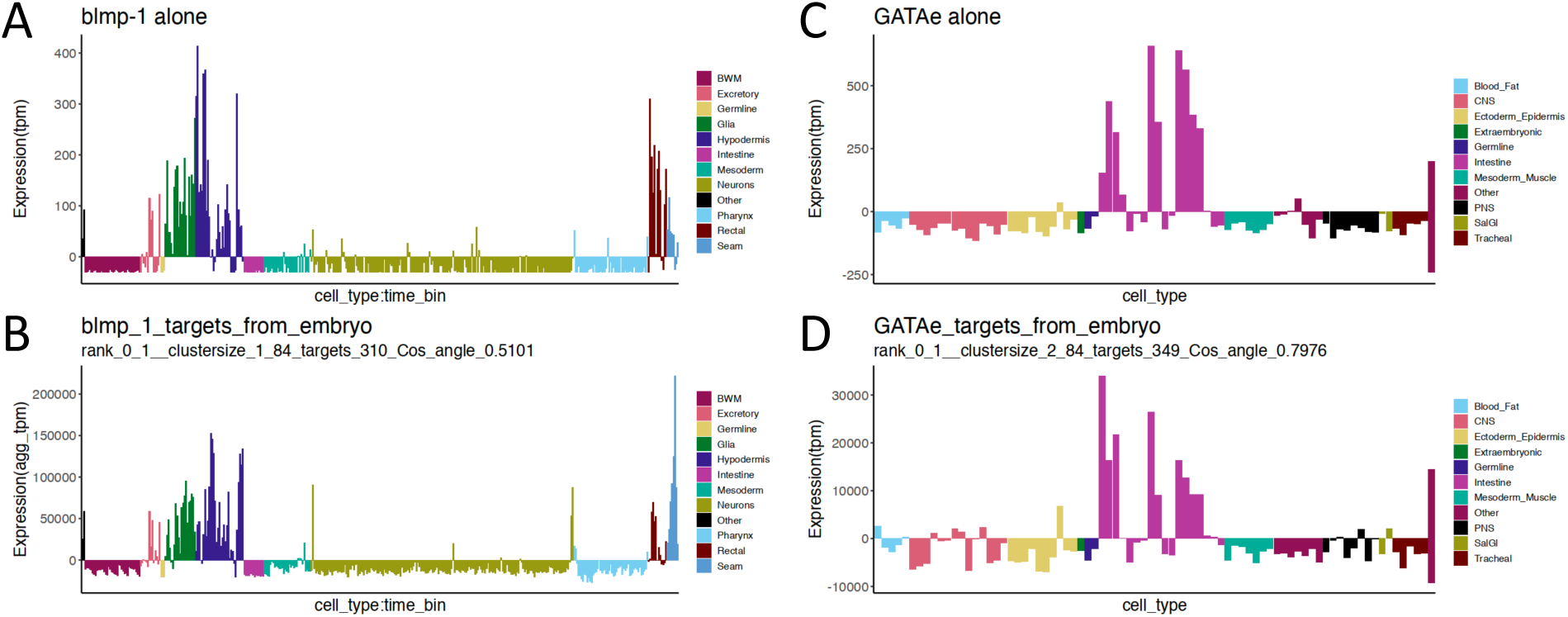
TF expression correlates with target gene expression. **A-D** Aggregate target expression reflects the TF expression for the worm *blmp-1* (A) TF and (B) targets, and the fly GATAe (C) TF, and (D) targets. Cell types are arranged along the x axis by broad cell class and sorted alphabetically within each class. The worm cell types are further divided into time bins (from Packer et al., 2019, Table_S7).

The aggregate expression patterns of these nonHOT targets of *blmp-1* and GATAe are remarkably similar to the patterns of the respective TFs. *blmp-1* in worms is expressed in a variety of cell classes, including hypodermis, seam cells, glia, excretory cells and rectal cells with the aggregate expression of the target genes closely resembling the pattern of the TF. GATAe in flies is expressed in several different cells of the intestinal tract, again with its target gene pattern closely reflecting that of the TF.

To evaluate the correlation of TFs more broadly with their cell types, we calculated the cosine of the angle between the expression vectors of the TF and its targets across the different cell states: the embryo TFs against the expression in both progenitor and terminal cell states, the L1, L2 and L3 larval stage TFs against expression in the L2 stage, and the L4 and young adult TFs against expression in the young adult stage (see **Methods**). For flies, we examined the embryos TFs against the embryo expression data. The TFs with the most highly correlated targets by this metric include many well-known TFs in both organisms as well as additional less-studied TFs (**Suppl. File 2**). Manual inspection of many TF target comparisons showed that, at a cosine of 0.2 or greater, the TF/target patterns still had readily recognizable similarity. In total, 142 worm TFs had an angle >= 0.2 in one or more of the four comparisons and 89 fly TFs had an angle >= 0.2 in embryos. Some of the cosine angles are negative, possibly signaling a repressive effect of the TF, with more TFs having more negative values in fly than worm. In worms, *zip-5* is highly expressed in intestine, yet its targets show negative values in the intestine as well as in most of the other cells with high *zip-5* expression. In flies, su(Hw), known to act as a repressor in many circumstances (Geyer and Corces 1992), shows negative values for the aggregate target expression in cell types in which su(Hw) itself is expressed.

To learn which of the targets was contributing to the aggregate signal, we compared each of the targets individually with its TF, with the idea that targets resembling the TF pattern might be the result of functional TF binding. We also compared an equal number of random targets to each TF pattern and compared the two distributions to estimate the likelihood of obtaining such a TF-target distribution by chance, using a wilcoxP test. The results (**Suppl. File 2**) suggest that most of the cosine angles greater than 0.2 for worms are significant. For flies the correspondence between the cosine value and the wilcoxP significance is less well correlated, but cosine values above 0.4 are almost all significant.

A limitation of this approach is that for targets expressed in multiple cell types under the control of different regulatory elements (and TFs), the expression pattern of the targets may be quite different from that of the TF. A simple illustration is provided by *hlh-1*, the worm myoD homolog, with a cosine angle of 0.42. Its targets are highly enriched in body wall muscle expression and the GLR mesodermal cell, where *hlh-1* is expressed. But the aggregate targets also show expression in other mesodermal cells, where *hlh-1* is not expressed (**Suppl. Figure 4 A**). Satisfyingly, *hlh-8*, a homolog of *hlh-1*, with a cosine angle of 0.39, is expressed specifically in these mesodermal cells and not in body wall muscle, and the aggregate targets of *hlh-8* show enrichment in both the mesodermal and body wall muscle (**Suppl. Figure 4 B**). In this case the expression of the TFs and their targets is simple enough that it doesn’t depress the cosine angle overly, but clearly for complicated situations very low values could result. Other issues might include TFs that are both positive and negative regulators, effectively canceling any signal. In both analyses, there were many TFs that failed to show a correlation between the TF expression pattern and the target patterns. The basis for these essentially negative results is unclear. Some experiments may simply have failed, despite passing quality metrics. Other TFs may have only a fine-tuning role, with effects too subtle to measure with these methods.

### Modeling gene expression patterns

We next investigated if the signal from the ChIP sites and their targets could be used to model gene expression. We chose to use a random forest, machine learning algorithm because it accommodates nonlinear relationships and provides a relative measure of the importance of the individual TFs in predicting expression. As input we used the peaks and their targets, with a row for each metapeak-target relationship (targets with multiple metapeaks had multiple rows) and a column for each TF, with values of the matrix being the relative peak strength of the TF binding event (the predictor matrix). The response variable was the cell type gene expression as measured in the scRNA-seq data, which does not distinguish between isoforms. The algorithm then produces an output matrix, giving the relative importance of each TF for determining the target values for each cell type independently. Running the model on multiple cell types produces an output matrix that gives the relative importance of each TF for determining the relative target values for each cell type and an estimate of the goodness of fit.

We modified the input matrix in various ways, testing more than 40 different models in the embryo terminal cell data. We used different filters for the metapeaks to be considered, modified the peak rank by distance from the TSS, considered only peaks from TFs expressed in the cell type above a threshold, and looked at alternate targets as well as primary targets (see **Methods**). Using the estimate of the goodness of fit produced by the algorithm, we compared the performance of the different models across all the cell types and in turn across the different life stages, selecting models that performed well across many cell types. The top model for worms considered only genes that were not targets of HOT sites, required the TF to be expressed in the cell type at least 5% of the maximum across all the cell types, used the peak rank modified by distance to weight proximal peaks more strongly, removed singleton peaks and used only primary or close alternate targets (see **Methods**). The relative importance of the TFs in each of the cell types defined in the single cell data in the worm embryo is illustrated in **Figure 7** for the model with the best fit across the samples. The model reproduced many known relationships, e.g., in the worm *elt-1* and *elt-3* were of high importance in the hypodermal cell types, *hlh-1* and *unc-120* in the body wall muscles, *elt-2* in the intestine and even *che-1* in the ASE neurons. But it also suggested the importance of other less well-characterized TFs in these and other cell types (**Suppl. Figure 5**).

**Figure 7.**
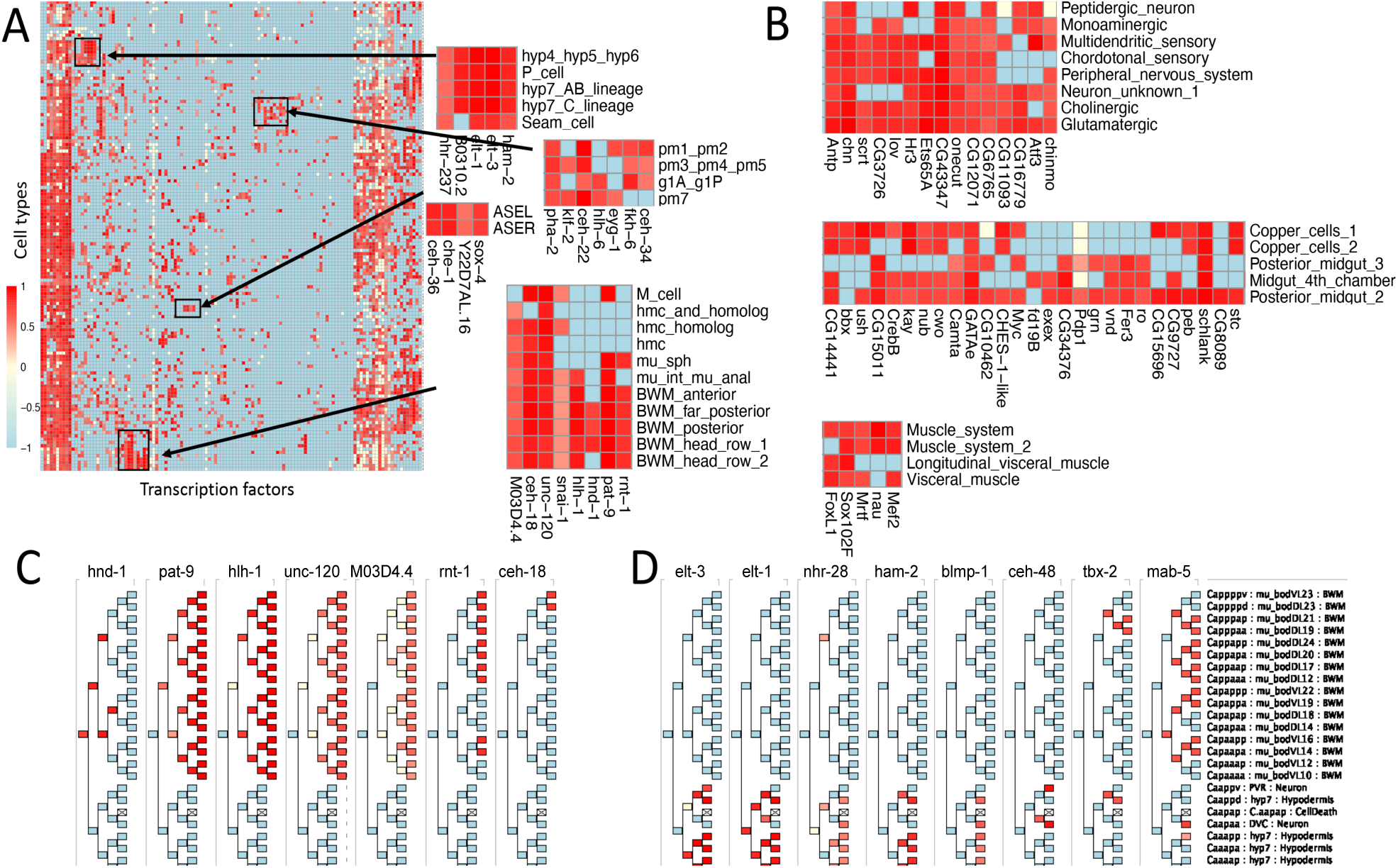
The relative importance of TFs in cell type gene expression. **A.** A heatmap of the relative importance of worm TFs in predicting gene expression in the embryo terminal cell types. The importance is indicated by the intensity of the red color from yellow (no importance) to dark red (most important). Light blue indicates that the factor was not expressed above threshold in that cell type. Clusters of TFs and cell types (black boxes) are blown up on the right, showing the detected relationships of well-studied and novel TFs and the cell types in which they are important. **B**. Clusters of fly TFs and cell types in which they are important, selected from the full heatmap in Supplemental Figure 5. Color scale as in A. **C**. **D.** TFs important in the Ca lineage. **C.** TFs important in the Cap lineage, which produces exclusively body wall muscle cells, and are arranged by onset of their importance. **D.** TFs important in the Caa lineage or in patterned expression in both lineages. The Caa lineage produces primarily hypodermal cells but also some neurons and cell deaths. The lineage names are given on the right, along with the specific cell type in the embryo. Body wall muscle cells are labeled mu_bod followed by letters to indicate Dorsal/Ventral, L/R and a number indicating the row of the cell, with 24 most posterior. Color scale as in A. Cell types not detected in the single cell data are indicated by an X.

We also ran the best model on the combined fly ChIP-seq and scRNA-seq data. Because of the low level, broad expression of many fly TFs (**Suppl. Figure 5 D**), we increased the threshold for minimal TF expression to be greater than the mean expression across the cell types plus one standard deviation. Again, the model captured the importance of TFs known to function in certain cell types, such as GATAe in the gut and the TFs involved in muscle. But overall the fly model appears less discriminating than the worm. Whether this lack of definition reflects the different biologies of the two species or results from limitations of the scRNA-seq or ChIP-seq data is unclear.

Examination of the worm lineage heatmap of the TF importance suggested that the model was capturing some of the dynamics of regulation during development. To facilitate the visualization of such relationships we developed a program that allows the user to view the importance of the various TFs in arbitrary subsets of the lineage and TFs. For example, in the Ca lineage (**Figure 7 C**), the TF *hnd-1* is important in the Ca cell itself, but only in the progeny that give rise to body wall muscle. As *hnd-1* importance wanes, *hlh-1* and *pat-*9 become increasingly important, followed by *unc-120* and *M03D4.4*, each with lesser importance. In the Caa branch, which gives rise predominantly to hypodermis, *elt-1* and then *elt-3* become important, with *nhr-28* later and of lesser importance (**Figure 7 D**). The *ceh-48* TF gene, a known regulator of neural development, only becomes important once the lineage becomes restricted to producing neurons. The restriction of TF expression clearly limits where TFs can be considered important, but the ChIP-seq binding data are central to determining the importance of each factor during development. The program is available as a java jar file for users to install on their own computers to explore TF importance in any of the lineages.

### Data access

All of the data have been deposited at the ENCODE Data Coordinating Center, with the exception of experiments concluded after October, 2022, when the Center stopped accepting submissions. To host these more recent data, and to provide a user-friendly interface, we have developed a website that provides access to all the data sets associated with each experiment, including the non-TFs that were not analyzed further in this report. This website also serves links to UCSC browser displays and has other features. It provides a file of all the peaks with associated metadata for all the TF experiments, such as the target and its distance from the peak, the position of the peak relative to the gene, etc. Interactive heat maps of the importance of each TF in predicting the gene expression pattern of each cell are available as html files. Finally, users can also access the Java program that displays the temporal importance of factors in the embryonic lineage of the worm.

## DISCUSSION

We have described here the results of a long-term project to discover the binding sites of the TFs in two key model organisms, *Caenorhabditis elegans* and *Drosophila melanogaster*. The effort has produced ChIP-seq binding profiles for more than 90% of the fly TFs and more than half of targeted TFs in worm. Most of these TFs have been assayed at the stage where the TF is maximally (or near maximally) expressed but some have been assayed at multiple stages, with a total of 674 experiments in the fly and 519 in the worm.

Combined, these experiments define 3.6 M peaks in the fly and 0.9 M peaks in the worm. The peaks often overlap in genomic location, creating what we have termed metapeaks, with each metapeak containing anywhere from a single peak to peaks from hundreds of TFs. Together these metapeaks appear to sample much of the regulatory DNA of each genome, based on the overlap with ATAC-seq data and inspection of defined regulatory regions in each species.

The fly and worm datasets have remarkable similarities, but one clear difference is the number of peaks found in each experiment and the number of peaks associated with each gene. Combined with the larger number of TFs assayed in the fly, the fly has about four times as many peaks detected as the worm. This increase is not reflected in a similarly increased number of metapeaks (only about a 50% increase). While the amount of the genome covered by peak signals is about two-fold greater in fly (57 Mb vs 27.5 Mb), the fraction covered rises only about by half, from 27% to ∼40%. The result is a considerable increase in the number of peaks per metapeak. Greater cross-reactivity of the GFP-antibody with fly DNA binding proteins might be one explanation, but the pattern of peak signal strength vs metapeak occupancy is almost identical between the fly and worm, using similar fractional cutoffs in grouping. Whatever the reason, a practical consequence has been that more stringent relative thresholds in fly were useful in detecting TF-target site similarities and random forest modeling.

Using a straightforward distance metric, we have assigned each peak and metapeak to a target and a possible alternate target gene. Genes have widely varying numbers of associated peaks and metapeaks, with some genes having tens of regulatory regions as defined by the metapeaks. Those metapeaks with the largest number of peaks, or HOT sites, contain some of the strongest relative peak signals, are most closely located near TSSs and have target genes that are broadly expressed. Together these results suggest that HOT sites are important functional regions that likely contain very open chromatin where TFs in almost any cell type are able to bind. By contrast those metapeaks with fewer peaks (lower occupancy) appear to drive more tissue specific expression patterns. Even these sites can have large numbers of TF binding sites. Whether cell type specific sites of open chromatin act as similar sinks for TF binding is unclear from these whole animal datasets. Functional redundancy and TF cooperativity could also contribute to this high density of sites (Wu and Lai 2015).

Taking advantage of single cell RNA-seq data for each organism, we find a clear relationship between the expression pattern of a TF with its targets, presumably reflecting functional binding events. Further, we have used the ChIP binding sites to drive a random forest model of cell type specific regulatory relationships. This analysis identified a subset of TFs that contribute broadly across many cell types to regulate target gene expression. Most importantly, this analysis also identified many TFs with significant contributions to gene expression that is restricted both temporally and spatially. To highlight this specificity, the impact of distinct TFs on gene expression can be distinguished between parent and daughter cells in the embryonic lineage of the worm, as well as for a single neuron pair. In many cases, TFs known to have a role in a specific cell type show similar patterns and scores with TFs that have not yet been linked functionally to that cell type, suggesting worthwhile avenues of investigation into the relationship of these TFs in development or activity of that cell type. The RNA-seq was key in limiting the cell types the TFs might be active, but the ChIP-data itself determined the importance of each TF in generating the pattern of expression. Although this analysis does not identify individual target genes that depend on a given TF, it increases the overall resolution of the whole animal ChIP-seq data, as well as its relevance for further investigation into TF function.

Beyond the ChIP-seq data, the GFP tagged strains present a valuable resource to the community. We have produced tagged strains for 508 TFs in the worm and 616 tagged strains in the fly along with additional strains for other genes. These strains have already been highly used, with the stock centers reporting requests for 17,293 and 4,334 strains for the fly and worm respectively.

The resource certainly has its limitations. The data comes from whole animals, mostly from a single stage. TFs expressed in only a single cell type where that cell type is represented by only a few cells in the animal seem unlikely to produce robust ChIP signals. Indeed, in the worm, we stopped performing ChIP for strains with very limited expression, because of challenges in getting passing samples with meaningful peaks. Nonetheless it is worth noting that the TF *che-1*, only expressed in the ASE neurons (two cells of that type in the 959 somatic cells of the worm) and known to be a driver of its differentiation (Patel and Hobert 2017), was detected as one of the most meaningful TFs in that cell type in the random forest modeling. In contrast, in the experiments with *mec-3*, expressed predominantly in four touch cells in the embryo, the targets do not include known genes that are not expressed in *mec-3* mutants (Etchberger et al. 2009). Further, TFs may be redeployed driving different genes even in different cell types at different stages in development and by only assaying one stage, we have missed these events. However, in aggregate, this resource provides hundreds of worm and fly strains, and millions of in vivo binding events that are already heavily utilized by the model organism community. With increasingly sophisticated computational approaches, we expect that these data will be critical to investigating and understanding complex gene regulatory programs during development.

## ACKNOWLEDGEMENTS

The authors thank: the ENCODE Data Coordinating Center for accepting and providing data access; the Yale Center for Genome Analysis (YCGA), especially Bryan Szewczyk and Irina Tikhonova for library preparation and Christopher Castaldi for sequencing (the YCGA is funded by the National Institute of General Medical Sciences of the National Institutes of Health under Award Number 1S10OD030363-01A1); the Bloomington Drosophila Stock Center (BDSC) for distributing the fly GFP-tagged TF strains; and the Caenorhabditis Genetics Center (CGC) for distributing the worm GFP-tagged TF strains (the BDSC and CGC are funded by the National Institutes of Health (NIH) Office of Research Infrastructure Programs P40 OD-018537 and P40 OD-010440, respectively). We would also like to thank WormBase (https://wormbase.org/) and the UCSC genome browser (http://genome.ucsc.edu) for displaying our data; the University of Chicago DNA Sequencing Facility (DNASEQ) for sequencing recombineered BACs (the DNASEQ is supported by the National Cancer Institute, Comprehensive Cancer Center grant P30CA014599); Donald Court for sharing his recombineering plasmids; Krishna S Ghanta and Craig Mello for providing CRISPR technical advice. This work was supported by NIH grants U41HG007355 (R.H.W.) and NIH grant R01GM76655 (S.E.C)

## AUTHOR CONTRIBUTIONS

MK performed worm ChIP-seq, oversaw worm sample processing and sequencing of both fly and worm, and wrote the paper.

LG analyzed the data, created the website and associated tools, and wrote the paper.

AV performed worm ChIP-seq and managed fly sample processing.

BCL performed fly ChIP and managed fly sample processing.

JG performed data analysis.

JX performed data analysis.

SS performed worm culture.

EF generated

GFP-tagged

TF worm strains.

ATP generated

GFP-tagged

TF worm strains.

CH oversaw worm strain production and edited the paper.

DV generated

GFP-tagged worm strains.

AH generated

GFP-tagged

TF fly strains.

WF generated

GFP-tagged

TF fly strains.

MW performed fly ChIP.

GW performed fly ChIP.

VH performed fly ChIP.

ZL performed fly ChIP.

MK performed fly ChIP.

KPW oversaw the fly ChIP.

RA oversaw the fly ChIP.

MG performed data analysis.

LDH performed data analysis.

SEC coordinated fly strain generation and wrote the paper.

VR coordinated the worm ChIP-seq project and wrote the paper.

RHW was the lead coordinator of the entire project, analyzed the data and wrote the paper.

## STAR METHODS

### Strain generation and resource

For both fly and worm we revised our lists of transcription factors (TFs), removing those that WormBase or FlyBase had designated as pseudogenes, and reclassifying others after consulting recent publications (Ma et al. 2021; Narasimhan et al. 2015) and considering our own results with factor subcellular localization in transgenic lines. Our revised list contains 658 fly TFs and 833 worm TFs (**Suppl. File 1**).

For both species we generated strains with individual TFs tagged with Green Fluorescent Protein (GFP), but the details differed, taking advantage of the different resources available.

For the fly, C-terminally GFP tagged BACs (bacterial artificial chromosomes) were used to create transgenic lines as described previously (Kudron et al. 2018; Venken et al. 2009). In brief, GFP was recombineered into the smallest P[acman] BAC that also contained either a predicted insulator or a portion of the next nearest gene upstream of the TF. GFP tagged BACs were integrated into specific attP sites through injection using the phiC31 integration system. Some additional GFP BACs were sent for injection and balancing at Genetivision (https://www.genetivision.com/). In general, these BACs were either injected into parent strains VK31 or VK37 such that the TF-GFP would integrate into a different chromosome than the endogenous TF. Homozygous lines were able to be generated for ∼90% of the TF-GFP lines. Altogether, tagged strains are available for 616 TFs. Lines were also constructed for an additional 56 nonTF genes (**Suppl. File 1**). All lines are available from the BDSC (https://bdsc.indiana.edu/stocks/misc/taggedtnx.html).

For the worm for most of the project, GFP-tagged fosmids from the modENCODE TransgeneOme resource (Sarov et al. 2012) were introduced by bombardment (Kudron et al. 2018). Bombardment resulted in low number of copies of the fosmid (from 1 -10) at undetermined sites in the genome. However, not all TFs were represented in the library and by 2017 the remaining available fosmids had repeatedly failed to yield GFP-tagged transformants. The development of CRISPR methods for the worm provided an alternative, and also offered the chance to generate strains in which the endogenous copy of the gene is tagged, avoiding possible issues of higher copy numbers, competition with the endogenous copy and genome position effects. Over the course of the project we used three different methods. In early experiments we used the SapTrap method (M. L. Schwartz and Jorgensen 2016) (strains RW12167-RW12175) but then switched to using Roller plasmid as a co-injection marker (Dokshin et al. 2018; Ghanta, Ishidate, and Mello 2021). We targeted C-terminal fusions but in the rare cases where these failed, we tagged the N-terminal end. Roller progeny were selected and their progeny evaluated for GFP expression with fluorescence microscopy. We used this method for the bulk of CRISPR strain collection (strains RW12178-RW12346). One issue we encountered with the method was the generation of successfully GFP-tagged strains which also exhibited the Roller phenotype and the two phenotypes co-segregated. To avoid this problem and to use all commercially generated reagents, toward the end of the project we switched to using a *dpy-10* repair template as a co-injection marker (Paix, Folkmann, and Seydoux 2017; Paix et al. 2015). This method generally yielded hundreds of Roller progeny per 20 injected worms, with some injected worms yielding as many as 100 Roller progeny. Interestingly, these “jackpot” plates often had Dpy progeny, indicating that both copies of the embryo genome had been edited. These Dpy animals frequently gave rise to Dpy and non-Dpy progeny, confirming that the second edit had been post-zygotic (Farboud, Severson, and Meyer 2019). To avoid possible problems with integration events not producing detectable fluorescent signal, we tested up to 100 progeny of Roller animals from “jackpot” plates. Overall, this method seemed to provide the greatest success (strains RW12347 and after).

Because of the potential differences between the strains generated by fosmid bombardment and CRISPR editing, we generated CRISPR lines for TFs that had also been tagged using fosmid bombardment and compared the GFP expression and ChIP-seq results for both strains in detail (**Suppl. Table 1**). We tested factors at different stages and tissues of expression, looking at the success as measured by IDR, the number of binding sites detected and the intensity of GFP signal (expression) in the paired strains. We saw no consistent differences.

Altogether, tagged lines are available for 508 TFs and another 45 lines were constructed for nonTF genes. Some of these were contributed by the wider community (see **Suppl. File 1** for details.) Almost all strains are available from the Caenorhabditis Genetics Center (CGC).

### Fly growth and ChIP

As described previously, transgenic flies were expanded in vials and bottles containing standard Drosophila media at ∼22℃. Larvae, white pre-pupae(WPP) and adults were collected from these expansion bottles. Pupae were obtained by collecting WPP and incubating for 1-2 days at ∼22℃. For embryonic collections, 0-8 day old adult flies were transferred to embryo cages capped with apple juice plates. After a 24 hour preincubation, apple juice plates were replaced at appropriate intervals, restricting the collected embryos to their desired developmental stages. For late embryonic stages, collection plates were removed and maintained at room temperature prior to chromatin collection.

Chromatin was collected and immunoprecipitated (IP) using the goat anti-GFP described previously (Zhong et al. 2010; Niu et al. 2011; Kasper et al. 2014; Kudron et al. 2018). However, due to its limited supply, later experiments used anti GFP Ab290 (Abcam) (**Suppl. File 1**). In total, ChIP was successfully performed for 677 (604 different TFs) experiments with TFs and another 63 experiments (56 different genes) for nonTFs. Of the 12 tagged TF strains without ChIP, six failed to yield data and for another six, ChIP was not attempted.

### Worm growth and ChIP

For the modERN project, worm growth and ChIP were conducted as previously described (Zhong et al. 2010; Niu et al. 2011; Kasper et al. 2014; Kudron et al. 2018) with the following modifications. Embryonic stages were collected following bleaching and resuspension in M9 at 20°C with shaking until the desired embryonic stage was visualized by DIC microscopy. Larval and the young adult stages were synchronized by bleaching and L1 starvation followed by plating the arrested L1s on peptone-enriched plates seeded with OP50 bacteria. Animals were grown for 6 hrs at 20°C for L1 collection, or longer to achieve the desired stage using DIC microscopy (Brenner 1974). GFP and DIC images were recorded immediately before collection using a Zeiss Axioplan 2 microscope. Worm samples were then cross-linked with 2% formaldehyde for 30 min at room temperature and quenched with 1M Tris pH 7.5. The pelleted worms were rinsed with M9 buffer twice and then with 150 mM NaCl FA buffer (50mM HEPES/KOH pH7.5, 1mM EDTA, 1% Triton X-100, 0.1% sodium deoxycholate, 150mM NaCl) with one Roche Complete tablet (Catalog number 11697498001) before the pellet was flash frozen in liquid nitrogen and stored at -80°C. Worm pellets resuspended in a total volume of 1.5 mL of 150 mM FA Buffer containing one Roche Complete tablet, 25 ul 1M DTT and 125 ul of 100 mM PMSF per 25 mL chilled FA were sonicated using a microtip to achieve mostly 200–800 bp DNA fragments.

Worm lysate was measured by Bradford assay. Lysate corresponding to 2.2 or 4.4 mg was resuspended in 150 mM FA Buffer with Complete protease inhibitors along with 1:20 20% N-lauroylsarcosine sodium salt and spun for 5 minutes at 13,000 g at 4℃ in a non-stick Eppendorf tube. Samples were transferred to a fresh tube and 10 percent of the lysate was removed as input. Input samples were stored at -20℃ until the following day. ChIP samples were immunoprecipitated using 7.5 μg anti-GFP antibodies (gifts of Tony Hyman and Kevin White) overnight at 4℃ with rotation.

The following day, input samples were thawed and 2μl of 10 mg/mL RNase A was added and samples were digested at room temperature for 2 hours. Then input samples were brought up to 300 μl using Elution Buffer (1% SDS in TE pH 8.0 with 250 mM NaCl), and 4 μl of 10 mg/mL Proteinase K (Roche 03115887001) was added and they were incubated at 55℃ for 4 hours.

In the meantime, the ChIP samples were incubated with protein G sepharose beads for 2 hours at 4℃ with rocking (GE Healthcare Cat. # 17-0618-01). Beads were first washed 4 times in 150 mM FA buffer. All subsequent spins were at 2500 x g for 2 minutes each. The beads were washed 2 times with room temperature 150 mM FA buffer for 5 minutes and then 1 time with 1M NaCl FA (50mM HEPES/KOH pH7.5, 1mM EDTA, 1% Triton X-100, 0.1% sodium deoxycholate, 1M NaCl) for 5 minutes. Samples were then transferred to new tubes and washed one time with 500 mM NaCl FA (50mM HEPES/KOH pH7.5, 1mM EDTA, 1% Triton X-100, 0.1% sodium deoxycholate, 500mM NaCl) for 10 minutes, 1 time with TEL buffer (0.25M LiCl, 1% NP40, 1% sodium deoxycholate, 1mM EDTA, 10mM Tris-HCl, pH8.0) for 10 minutes and then 2 times with TE pH 8.0 for 5 minutes. In order to elute the immunocomplexes, the beads were incubated at 65 ℃ two times for 15 minutes each with 150 μl Elution Buffer with gentle vortexing every 5 minutes. The two elutions were pooled after spinning down and 1 ul of 20 mg/mL Proteinase K was added to each ChIP sample and the tubes were incubated for one hour at 55℃. Both the eluted ChIP samples and the input samples were then incubated overnight at 65℃ to reverse crosslinks. All samples were purified using either a minElute PCR Purification Kit from Qiagen (Catalog number 28006) or a ChIP DNA Clean and concentrator kit from Zymo (ZD5205).

In total, ChIP was successfully performed for 508 (350 different TFs) experiments with TFs and another 65 experiments (45 different genes) for nonTFs. Of the 149 tagged TF strains without ChIP, 19 failed to yield data despite repeated attempts, 39 were deemed to be poorly expressing, 11 were deemed orphan nuclear hormone receptors and set aside, 4 had a phenotype and for another 20s ChIP are ongoing.

### Worm and fly ChIP-seq library prep and sequencing

ChIP and input control samples (genomic DNA from the same sample) for two technical replicates for worm or three for fly were used for library preparation and sequencing as previously described for the modENCODE and modERN projects (Zhong et al. 2010; Nègre et al. 2011; Kasper et al. 2014; Kudron et al. 2018). More recently, samples were processed at the Yale Center for Genomic Analysis (YCGA), using the following methods (**Suppl. File 1**): Libraries were generated using approximately 5-10ng of DNA that were end-repaired, A-tailed, adapter ligated and PCR enriched (8 -10 cycles) using the KAPA Hyper Library Preparation kit (KAPA Biosystems, Part#KK8504). Indexed libraries were quantified by qPCR using the Library Quantification Kit (KAPA Biosystems, Part#KK4854) and inserts were size distributed using the Caliper LabChip GX system. Samples with a yield of ≥0.5 ng/ul were used for sequencing.

Sample concentrations were normalized to 10 nM and loaded onto an Illumina NovaSeq flow cell at a concentration that yields > 5 million reads passing filter clusters per sample. Samples are sequenced using 100bp paired-end sequencing on an Illumina HiSeq NovaSeq Sequencer according to Illumina protocols. A positive control (prepared bacteriophage Phi X library) provided by Illumina is spiked into every lane at a concentration of 0.3% to monitor sequencing quality in real time. De-multiplexed sequencing data passing internal quality controls at YCGA were transferred to the University of Washington for downstream processing.

### Peak calling

During the course of the project sequencing methods evolved, the SPP peak caller went through multiple rounds of improvement and our internal pipeline changed (Kudron et al. 2018). To create a uniformly called set of peaks for the analyses presented here we processed all the ChIP-seq experiments through the ENCODE Transcription Factor and Histone ChIP-Seq processing pipeline version v1.3.6, https://github.com/ENCODE-DCC/chip-seq-pipeline2/tree/master. (This version of the processing pipeline was used for all data submitted to the ENCODE DCC as well as data generated after October 2022.) In all cases the bwa program was used for alignment (Heng Li and Durbin 2009; Heng Li 2013). The version of bwa in that pipeline is 0.7.17 https://bio-bwa.sourceforge.net/. For very early experiments that generated only short single ended sequence reads, the alignments done with the earlier version of bwa. 0.7.8 were used. The peak caller used for all experiments is SPP version 1.15.5 (https://hbctraining.github.io/Intro-to-ChIPseq/lessons/peak_calling_spp.html) <u>(Kharchenko, Tolstorukov, and Park 2008).</u> Because the ENCODE DCC ceased to accept data in October, 2022, these new peaks calls are not available there but are available only through the website https://epic.gs.washington.edu/modERNresource and also through the SRA.

### Clustering of transcription factor peaks into metapeaks

Exploiting the similarity between TF peaks and sequence reads aligned to the genome, we aggregated TF peaks across all the ChIP-seq experiments for each species and called metapeaks. These called metapeaks represent the bounds of the clusters of TF peaks. The TF peaks are then assigned to metapeaks based on proximity and overlap with the metapeaks.

We first aggregated all the peaks from the TF experiments for each species into a single bed file. SPP sometimes called several peaks from a single experiment that overlapped exactly with the exact same genomic span but different apices. These exactly overlapping peaks were trimmed, so that they were contiguous and non-overlapping. The endpoints of these trimmed peaks were the midpoint between apices. The apices were unchanged.

We next formed a signal track of all the aggregated trimmed TF peaks as a bedGraph file, as described at http://genome.ucsc.edu/goldenPath/help/bedgraph.html, using the program bedtools genomecov -bg (Version: v2.29.0) https://bedtools.readthedocs.io/en/latest/content/tools/genomecov.html.

The MACS2 program (version 2.2.4) https://hbctraining.github.io/Intro-to-ChIPseq/lessons/05_peak_calling_macs.html was selected to call the metapeaks from the aggregated TF peaks. Unlike SPP, the MACS2 program can be run without a control signal track, and for the TF peaks there is no control signal track.

The bdgpeakcall subcommand of MACS2 was used to call the metapeaks from bedGraph files. Inspection of the results showed that clusters of peaks could sometimes overlap on their edges and depending on the extent of the overlap the valley between the two putative metapeaks could be of differing depths. To split these overlapping metapeaks, we iterated the peak calling procedure using a minimum peak length of 50 bases with cutoff thresholds from 2 to 50. This created 49 output files in narrowPeak bed format https://genome.ucsc.edu/FAQ/FAQformat.html#format12. As the cutoff increases, metapeaks with an internal valley will split into 2 metapeaks when the cutoff exceeds the minimum of the valleys. Using a range of cutoff values makes it possible to split these broad metapeaks with internal valleys into separate metapeaks.

To come up with a final set of metapeaks, the multiple cutoff metapeaks were arranged into a tree structure. The root of the trees was the metapeak called with cutoff 2. The children metapeaks were the metapeaks from next higher cutoff that overlap the lower cutoff metapeak. If the higher cutoff did not go above an internal valley, there was only one child, but if the higher cutoff goes above a single internal valley, there were two children. The tree was extended to the maximum threshold. Branches extending only one generation were trimmed to avoid over splitting. For the trimmed trees, each leaf metapeak was traced up the tree to when it was first created by a split of a metapeak, that is, it has a sister, and it is added to the list of metapeaks.

After all the leaf metapeaks were traced up to their origins, the genomic width of the selected metapeaks is expanded to fill the entire width of the metapeaks at root level of cutoff 2. This process is a compromise between splitting broad metapeaks with deep internal valleys and not splitting those metapeaks with very shallow internal valleys (depth of just one). The resulting metapeaks are also non-overlapping.

Each TF peak was then assigned to a metapeak as follows: If the TF peak apex was within the range of a metapeak, it was assigned to that metapeak. If the TF peak apex failed to overlap the range of a metapeak, but the peak region overlapped a single metapeak, it was assigned to that metapeak. If the TF peak region overlapped more than one metapeak range, it was assigned to the metapeak whose end was closest to the apex of the TF peak. If the TF peak did not overlap any metapeaks, then a new singleton metapeak was created for that TF peak.

### Conservation scores

The *D. melanogaster* (dm6) and *D. virilis* (droVir3) assemblies were aligned using blastz (Chiaromonte, Yap, and Miller 2002; S. Schwartz et al. 2003) using blastz parameters and scoring matrices recommended on the UCSC browser (Kent et al. 2002). Similarly, the *C. elegans* (ce11) and *C. remane*i (caeRem4) genomes were aligned. The alignments were chained using axtChain (Kent et al. 2003) and processed into nets by the chainNet and netSyntenic tools (Kent et al. 2003). Conservation scores were calculated by giving matching bases a score of 2, gaps a score of 0, and substitutions the following scores A/C=T/G=A/T=C/G=-2, A/G=T/C=-1. Scores were summed across the bases in the feature of interest.

### HOT site determination

In *C. elegans*, Araya et al., (Araya et al. 2014) defined thresholds for identifying high occupancy target (HOT) sites and extreme occupancy target regions (XOT) by creating an equivalent number of random binding regions with a size distribution matching the true set of binding regions in each of their developmental stages. Running this procedure 1,000 times on each stage, they determined the cutoffs at which fewer than 5% and 1% of the simulated binding regions have higher occupancies. We reproduced their procedure using our data and compared the resulting cutoffs with those using the 5% and 1% raw cutoff thresholds. The number of clusters identified at the 5% cutoff were similar using the raw and simulated thresholds, but more clusters were identified as ultra-HOT at the 1% threshold using the threshold based on the simulated data.

In an alternative approach, we calculated a HOTness score using a kernel density estimation (KDE) approach (L. Ma and A. Victorsen, unpublished) looking for regions of maximum point pattern densities as used, for example, in defining areas of geographical regions of high crime rates. The following parameters were used: bandwidth 300, cutoff score 0.1, cutoff peak 0.00001, and local peak height 30. Thresholds were defined by stage and by chromosome and across chromosomes within each stage using 1000 iterations randomly sampling from all observed peaks. This approach labels the largest number of peaks as HOT sites (**Suppl Figure 6 A**).

Other projects used thresholds based on the percentage of TFs studied that were bound in a given region. For example, in *Drosophila*, Moorman et al. (Moorman et al. 2006) defined those regions bound by 7 of their 7 transcription factors studied as HOT sites. In *C. elegans*, Gerstein et al. (Gerstein et al. 2010) defined those regions bound by 15 or more of their 22 factors as HOT sites. And in the human genome, Partridge et al. (Partridge et al. 2020) defined HOT sites as those regions occupied by one third of their assayed chromatin associated proteins. Given the more straightforward interpretation and reflection of the underlying data, the conservative estimate of HOT sites in comparison to other methods, as well as the large size of our data set, we chose to define our HOT site thresholds based on our own underlying data.

In *D. melanogaster* almost all samples were embryonic, but in *C. elegans* we experimented with setting HOT site thresholds per stage as suggested by Araya et al., (Araya et al. 2014) identifying the 1% and 5% raw threshold per stage as compared to identifying the thresholds for the data set as a whole. We found most sites were found in all of the developmental stages. Approximately 10% additional sites were identified as “HOT” when using the cutoffs set per stage rather than setting the cutoff across all stages with 92% of those were within a few percent of the cutoff. Thus, we chose to define the thresholds across the full data sets in both *C. elegans* and *D. melanogaster*, defining the 5% most highly occupied metapeaks as HOT sites and the 1% most highly occupied metapeaks as ultra-HOT sites (**Suppl Figure 6 B, C**).

### Overlap of metapeaks with chromatin features and ATAC peaks

#### D. melanogaster – ATAC-seq

Calderon et al., (Calderon et al. 2022) obtained 110,185 ATAC-seq regions from staged *D. melanogaster* embryos (11 overlapping time windows spanning the first 20 hours of development). As measured by an overlap of at least ten bases with ChIP metapeaks, by count 80% of ATAC regions are overlapped by a metapeak and 65% of metapeaks are overlapped by an ATAC region.

#### Chromatin state data - *D. melanogaster*

Kharchenko et al. (Kharchenko et al. 2011) identified 21,955 chromatin state regions (in S2-DRSC and ML-DmBG3-c2 cells using ChIP-chip), some of which were overlapping, and assigned the genome in 200 bp bins to one of nine chromatin states. We looked for overlap using just the apex of the metapeak, a region of 200 bases centered on the apex and the full metapeak, requiring an overlap of at least ten bases with the chromatin regions for the latter two measures. The three measures produced similar results, with the three states associated with actively transcribed among the most shared, and the states with heterochromatin showing less overlap (**Suppl. File 3**). The largest number of ChIP metapeaks and the largest number of metapeak bases overlap transcriptionally silent intergenic euchromatin, perhaps reflecting the wider range of genes active in the whole animal as opposed to cell lines.

#### C. elegans – ATAC-seq

Janes et al. (Jänes et al. 2018b) identified 42,245 elements accessible in at least one *C. elegans* stage. As measured by an overlap of at least ten bases, 27,052 of the ChIP metapeaks overlap 33,911 of Janes et al. elements (**Suppl. File 3**). In terms of overlap at the base pair level, in the embryonic data for example, 18% of bases labeled with a chromatin state overlapped an embryonic modERN metapeak and 98% of bases in embryonic metapeaks were overlapped by a region labeled with a chromatin state in the embryonic state data.

Janes et al. (Jänes et al. 2018b) further annotated their accessible regions defining 13,596 protein-coding promoters and 19, 231 putative enhancers. Review of the overlap between their regions and the metapeaks reveals the largest numbers of regions overlapped by count by the metapeaks are those labeled as enhancer and coding promoter. Metapeaks that overlap a an ATAC regions have larger numbers of peaks (30.8 +/- 60.0) compared to metapeaks not overlapping ATAC regions (2.7+/- 4.5) and, similarly, ATAC regions that overlap a metapeak have stronger signals (18.2+/-30.0) compared to those that don’t (5.0+/-5.4), suggesting that metapeaks with low occupancy are less likely to be observed in ATAC-seq data.

To examine further the 3,756 accessible regions annotated as an enhancer and that were not overlapped by a metapeak, we assigned those regions to target genes in the same way we assigned metapeaks to target genes. Those 3,756 accessible regions were found to be more distant from the transcription start site, typically more 3’ of the transcription start site, than were those accessible regions annotated by enhancer that did overlap a metapeak. Of those 2,910 genes targeted by enhancers (Jänes et al. 2018b) but not overlapped by a metapeak, 2,749 (95%) were targeted by other, non-overlapping metapeaks. Thus only 161 genes are identified as being targeted by enhancers (Jänes et al. 2018b) that are not targeted by a metapeak. Of those 161, there are only 80 that have a TPM of at least 100 in at least one cell type in the embryo (Packer et al. 2019).

#### *C. elegans* – eY1H comparison

Fuxman Bass et al (Fuxman Bass et al. 2016) examined interactions between 409 of *C. elegans* TFs and the 500 bases upstream of 3,125 of *C. elegans* gene TSSs, using an enhanced yeast one-hybrid assay (eY1H) to create a gene-centered physical protein DNA interactions (PDI) network. Their results contained 26,497 PDIs of which 21,714 were defined as high quality (dataset_EV1). These high quality interactions were between 2,576 target gene promoters and 366 TFs. They report finding overlap for 20% of the 46 shared TFs detected by eY1H and by ChIP as of the 2014 release of the *C. elegans* modENCODE project. For the current modERN dataset, there are 182 TFs shared by this project and the eY1H dataset. When looking in the 500 bases upstream of the TSS for the 14,198 TF/target pairs from those 182 TFs, 1213 (8.5%) of have at least one ChIP peak overlapping that region by at least 10 bases. Of the 11,756 high quality Fuxman Bass TF/target pairs, 945 (8.0%) overlap at least one ChIP peak.

To understand better the low level of overlap between the two data set, we examined the relationship between expression of the eY1H TFs and their targets in the single cell data sets, in the same way we had done for the ChIP-seq TFs and their targets. We compared all of the eY1H TFs against the embryo, L2 and YA scRNA-seq sets. Overall, the fraction of TFs with a cosine angle greater than 0.2 was lower in the eY1H data. For example, for the embryo, of 304 eY1H TFs with high quality targets, and more than one target, 8.8% (27/304) had a cosine angle > 0.2. (58 had only 1 target and no angle was calculated. The eY1H data set has only 123 have more than 10 targets; more than half the targets are associated with just 18 TFs). By contrast, 26.8% (33/123) of the ChIP embryo TFs had a cosine angle of > 0.2. Some notable TFs have low angles or very few targets in the eY1H set, e.g. HLH-1 has only one target. Of the eY1H that have a cosine angle of > 0.2, about half also have a high cosine angle in the ChIP data. Interestingly, those TFs have more than a third of TF-target pairs overlapping between eY1H and ChIP, with PHA-4 showing 54% overlap (20/37). For others, the eY1H data may be more informative data than the ChIP-seq data, e.g., ZTF-6 has a high angle (0.21) in the eY1H data and little overlap with the ChIP data (2/95) and a weak cosine angle (0.07) in the ChIP data.

### TF Pearson correlations for the worm and fly stages based on co-occurrence in same metapeak

Pearson correlations were used to evaluate how often TFs occur together in the set of metapeaks. Pairs of TFs that occur often in the same metapeaks will have higher correlation values. The initial set of metapeaks used in this analysis ranged in size of 2 peaks to 84 peaks, corresponding to the upper limit of non-HOT sites in the worm. A binary matrix was constructed for each life stage in the worm and fly. These matrices were metapeaks by TFs, with a value of zero indicating the TF is not in the metapeak and value one indicating there was at least one peak of that TF in the metapeak. For each lifestage in each species, if the number of TFs in the metapeak was less than two, that metapeak was not included in the calculations. For each of the binary matrices, a Pearson correlation was then calculated for each pair of TFs across the metapeaks. The correlation matrices were reordered by hierarchical clustering and displayed in heatmap form.

### Motif methods

For known motifs, we collected fly and worm motifs determined by *in vitro* experiments, including SELEX, PBM, and B1H, from the Cis-BP database (Weirauch et al. 2014). For the motifs of a TF, we concatenated all the unique position-weighted matrices (PWMs) into a single file in the MEME motif format. In total, 361 fly experiments and 118 worm experiments have known *in vitro* motifs. Locations of the motif occurrence in the genome were identified by FIMO (Grant, Bailey, and Noble 2011). We intersected the motif occurrence regions and the metapeaks regions by Bedtools (Quinlan and Hall 2010).

For motif inference, we first converted the ChIP-seq regions to fasta files using Bedtools (with genome versions dm6 and ce11 for fly and worm respectively). We then used STREME (v5.4.1) (Bailey 2021) with default parameters to infer TF binding motifs from those ChIP-seq regions. We compared the inferred motifs to known motifs by TOMTOM (Gupta et al. 2007). For each experiment, if any one of the best three inferred motifs matched any one of the known motifs for that TF, we counted it as an experiment with successful inference.

For each ChIP-seq experiment, we used each of the following criteria to create several different input subsets: (i) keeping all peaks with no filters; (ii) keeping top 20% peaks sorted by SPP score; (iii) removing peaks that fall in clusters with size larger than 277 or 85, respectively for fly and worm; (iv) removing peaks that fall in clusters with sizes larger than 53 or 31, respectively for fly and worm; (v) same as (iii), and further remove singleton peaks; (vi) same as (iv), and further removing singleton peaks. Note that STREME will report an error if there were too few or zero peaks after filtering, and such subsets of that experiment will be disregarded. To make fair comparisons, for each of the subset types from (ii) to (vi), we randomly sampled the same number of peaks from all peaks as well. We repeated the sampling three times and reported the average. All inferred motifs from all groups are available on https://github.com/Jiahao-Gao/TF_motif.

For the 311 and 397 experiments in fly and worm respectively where the TF did not have *in vitro* motifs, we compared their inferred motifs from group (i) to those from group (iv), described above. If any one of the top three inferred motifs from group (i) was significantly similar (TOMTOM q-value<0.05) to one of the top three inferred motifs from group (iv), we considered this experiment as one with consistent inference results.

### Peak and metapeak target assignment

We relied on the following assumptions to guide the peak and metapeak assignments to target genes. TFs operate at transcription start sites (TSS) and the closer the peak or metapeak is to the TSS, the more likely it is operating to influence gene expression of that target gene. The peak or metapeak can be either upstream or downstream to the TSS. It is not likely that a TF will influence gene expression of a target gene, if there is an intervening TSS of a different gene. It is ambiguous when a peak or metapeak lies between two different genes’ TSSs, and the distance to each of these two TSSs is not dramatically different. In such cases, more than one target gene assignment for a single peak or metapeak was made.

The following describes the algorithm used to assign peaks and metapeaks to target genes. Using the single base apex of the peak or metapeak as the location of a peak or metapeak, we found the closest TSS to the apex in either the positive or negative genomic direction and assigned the gene with the closest TSS as the primary target. Other information recorded about the assignment included the distance and direction to the TSS, whether the apex was within an exon or intron, or whether the apex was outside the gene, on either the 5’ or 3’ side of the gene. After assignment of a primary target, an alternate target was chosen if the gene with the next nearest TSS to the apex in the opposite genomic direction from the primary target was a different gene than the primary. The result was that an apex located between two genes was assigned both a primary and alternate target gene. The distance and other information recorded can be used to assess how confident the assignment is to the alternate target gene. If the distance is large compared to the distance to the primary target, the alternate target assignment may be less likely to be operational. In addition, the location of peaks and metapeaks relative to the features of the target genes and their transcripts were recorded, again using proximity of the peak to the feature. The recorded relationships included being extragenic (either 5’ or 3’) or intragenic (either an internal TSS’s, an exon or an intron). For genes with multiple TSS’s both the transcript and gene positions were recorded.

### Remapping fly single cell data

As the initial analysis of the single cell RNA-seq fly data set (Calderon et al. 2022) for TF-target relationships yielded disappointing results, we attempted to refine the annotation. To find additional cell types, we removed the very early cells (0-6 hours) along with yolk nuclei, subsetting their clusters 1, 5, 8, 9, 11, 12, 15, 16, 18, 21, 22, 23 to create a new cell data set (cds) (238,445 cells out of 547,805 starting cells). Reducing the dimensions of this subset using first 300 PCAs and then 2 dimensions with UMAP in monocle3 (https://cole-trapnell-lab.github.io/monocle3/) with default parameters produced a more highly structured map (**Suppl. Figure 7 A**). After reclustering and grouping some clusters, dividing others and incorporating the prior neural annotations (Calderon et al. 2022), we produced the anonymous annotation (**Suppl. Figure 7 B**). The cells had relatively low counts, and highly expressed genes that, based on other evidence, should have been very cell specific, had a low level background throughout the maps. These factors may have limited resolution and contributed to the need to use high expression thresholds in the random forest models. Calculating the TPMs for each gene in each cell type and weighting by the differential expression yielded a heuristic score for potential markers of each cell type. These markers were then compared to the fly in situ data set (https://insitu.fruitfly.org/cgi-bin/ex/insitu.pl) of the Berkeley Drosophila Genome Project and other literature to assign each cell type anatomical names (**Suppl. Figure 7 C, D**). After two additional rounds of refinement, we identified a total of 83 cell types in addition to the cell types removed from the cell subset (maternal, yolk nuclei, early germline, etc.) (**Suppl. File 4**). To obtain a more robust estimate of gene expression, we used a bootstrap calculation (1000 iterations with replacement) similar to that used by Packer et al. to obtain the TPM values of each gene in each cell type (Packer et al. 2019).

### TF versus target expression

To compare the expression of TFs with their targets, we used the median of the bootstrap values (Cao et al. 2017; Packer et al. 2019; Ghaddar et al. 2023) to generate a gene by cell type expression matrix. We then subtracted the mean value for each gene across cell types to reduce the impact of highly and broadly expressed genes (a mean centered matrix). The TPM values of the identified targets of the metapeaks containing TF binding sites were summed to generate an aggregate score and the cosine angle between the TF expression and the target expression was calculated in R using:

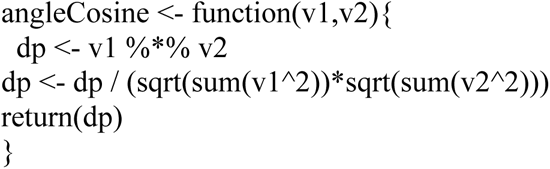

After exploring the impact of various methods of filtering the input peaks and targets, we settled on removing all targets of HOT sites and all singleton metapeaks. Removing targets of peaks with low relative signal strength did improve the scores of some TF-target pairs, but severely reduced the number of targets for some TFs. Expression plots were created in R using ggplot, assigning each cell type to a broad cell class and ordering the plots by the cell class and then the cell type.

### Random forest model of cell type expression

Regression models were trained to predict single cell gene expression in individual cell types from ChIP-seq TF binding sites. The goal of building these models was to determine which TFs were most important in determining expression in each individual cell type. A random forest machine learning method (https://urldefense.com/v3/ https://www.randomforestsrc.org/ <u>;!!K-Hz7m0Vt54!iUhkkDeBFzaT4erzyUP5WycP36QDAPYmgmzrL9D0w_nFOd2BH9xMRu8_P0d HwHJJM6qnAvdWl-uPHQ$</u>) was selected for this application because of the non-linear relationship expected between TF binding strength and gene expression. Also, the random forest model can be interrogated, after it is trained, for the most important TFs in predicting the gene expression in the cell type.

The independent variable in the predictor matrix is the TF binding signal strength. The binding signal is part of the output for each called peak by the SPP peak calling program. The dependent or response variable is each cell type’s gene expression profile. In order to relate the response variable to the predictor variables, each metapeak must be associated with one or more target genes whose single cell expression has been measured in the cell type. After modeling all the cell types in the embryonic, larval, and adult stages, a matrix of cell types by TF will be produced for each of those life stages. The values in this matrix reflect the relative importance of the TF in determining the expression of the target genes in the cell type.

Relating TF peaks to target genes is done on the basis of genomic proximity of the containing metapeak to the TSS. All of the TF binding sites have been grouped into metapeaks as described above. These metapeaks are assigned a primary target gene, which is the gene with a TSS closest to the apex of the metapeak. An alternative target gene may also be assigned to the metapeak, which is a gene in the opposite genomic direction from the primary target that has the closest TSS. Each binding site in the metapeak is assigned to the target genes assigned to the metapeak.

Multiple different models were built for all the cell types, depending on which metapeaks, TFs, and targets are selected for modeling, and any transformation performed on the signal binding strength of each binding site. Many of the ChIP-seq binding sites were determined in a single life stage, while some were done at multiple stages. When modeling the expression in a cell type, only binding sites measured in a close life stage were used in the modeling. Thus, embryonic ChIP-seq experiments were used to model embryonic single cell expression, while L1, L2, and L3 stage ChIP-seq experiments were used to model L2 larval expression. and adult and L4 ChIP-seq experiments were used to model adult expression for the worm. Only embryonic ChIP-seq experiments in the fly were used as this was the only stage for which single cell expression data was available.

The SPP peak calling program outputs a signal strength for each peak called. In order for these signal strengths to be used in the modeling, they need to be normalized so that each experiment is comparable. To do the normalization, the peaks in each experiment are sorted by their signal strength. They are then assigned a normalized signal strength between zero and one based on their rank in the experiment. The peak with the strongest signal in the experiment will be assigned a normalized signal of one, and the peak with the lowest signal strength in the experiment will get a zero normalized signal strength.

The predictor matrix is constructed with the independent variables in the columns, all the TFs measured in the experiments done in the life stage of the cell type being modeled. The rows of the predictor matrix represent the data points, an association of a cluster to a target gene. The values in the matrix in a given row are a measure of the binding site for each TF in the cluster. If a cluster is associated with more than one target, there will be more than one row in the matrix corresponding to that cluster. Genes that have no associated cluster are not represented in the model. Some genes will have multiple associated metapeaks, and there will be a row for the gene in the predictor matrix for each associated metapeak. The response vector is the single cell expression in the cell type being modeled for the target genes in the rows of the predictor matrix. The number of rows in the predictor matrix will equal the number of entries in the response vector.

Forty different predictor matrices and response vectors were constructed for each worm cell type, depending on which TFs were selected (feature selection), which metapeaks were selected, which targets were selected, and which function of normalized signal strength was selected.

### Transcription factor selection

1) only factors with nonzero measured single cell expression in the cell type
2) only factors with expression in the cell type of at least 5% of the maximum expression of all the cell types in the life stage.

### Metapeak selection

1) no HOT site and no singleton clusters, only those clusters with 2 to 84 peaks (of all life stages) in the cluster
2) clusters with 2 to 30 peaks
3) clusters with 2 to 84 peaks, but within 2KB of the target gene’s TSS or in the first intron or first exon of the target gene
4) clusters with 2 to 30 peaks, but within 2KB of the target gene’s TSS or in the first intron or first exon of the target gene
5) clusters with target genes that do not have a HOT site associated with them

### Metapeak target selection

1) Use primary targets only
2) In addition to primary targets, use alternative targets that are less than twice the distance from the cluster as the primary target

### Signal Strength

1) use the normalized signal rank without modification
2) divide the normalized signal rank by the log of the distance between the cluster and its target

Each cell type in each life stage were modeled with all forty different predictor models. When training a random forest, we used a procedure referred to as bagging, which means each decision tree is built with a random selection of the data points. The result is in an out of bag ensemble of data points, not used in the training, that can be used to assess the accuracy of each of the models. This out of bag ensemble also makes it possible to assess the importance of each TF in the accuracy of the prediction model by seeing how much the model error changes when a given TF is permuted in the trained model. Those TFs that cause the greatest increase in the model error when permuted, are the most important for the prediction.

The models were ranked by the root mean square error for each cell type. The model ranked best for the most cell types was selected as the best modeling approach for each life stage. For the worm, the model that used TFs with at least 5% of the maximum expression, no HOT sites and no singleton clusters, close alternative targets, and distance modified signal strength was judged to be the best.

For the fly embryonic stage, a different model performed best. This model used TFs where the expression was greater than the mean expression plus one standard deviation. Metapeaks more than 84 peaks were not included in the fly model, consistent with the worm models.

## SUPPLEMENTAL INFORMATION

**Supplemental Table 1.** Comparison of CRISPR vs bombardment method of worm strain generation on TF expression patterns and ChIP-seq results.

**Supplemental File 1.** Master spreadsheet of TFs for fly and worm. This list includes strain information, TF classification, ChIP-seq experiment data, including life stage, as well as information about non-TFs.

**Supplemental File 2.** Cosine angles for fly and worm TF-target gene relationships.

**Supplemental File 3.** Comparison of overlap between TF binding sites and chromatin accessibility.

**Supplemental File 4.** Re-annotation of embryonic fly cell types in scRNA-seq data.

---

**Supplemental Figure 1.**
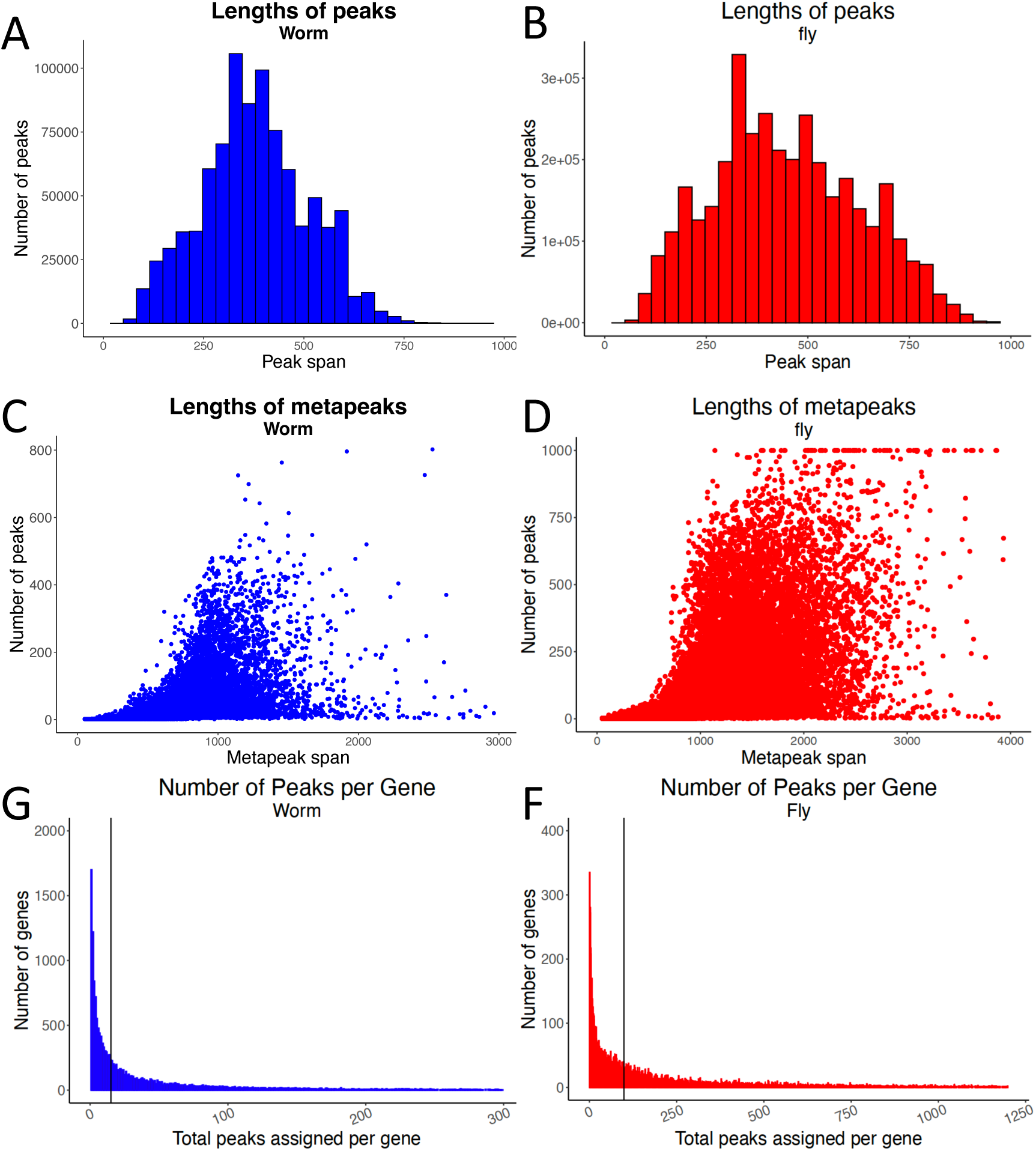
Peak features. **A, B**. The distribution of individual peak lengths in both worm and fly. **C, D**. The span of metapeaks increases only slightly with increased metapeak occupancy in worm (left) and fly (right), with almost all metapeaks less than 2 kb in the worm and less than 3 kb in the fly. **E, F** The number of peaks associated with each gene varies widely, with most genes having relatively few associated peaks, but with many with large numbers of associated peaks. The median number (vertical lines) of peaks per gene in worm was 15 and in fly 99.

**Supplemental Figure 2.**
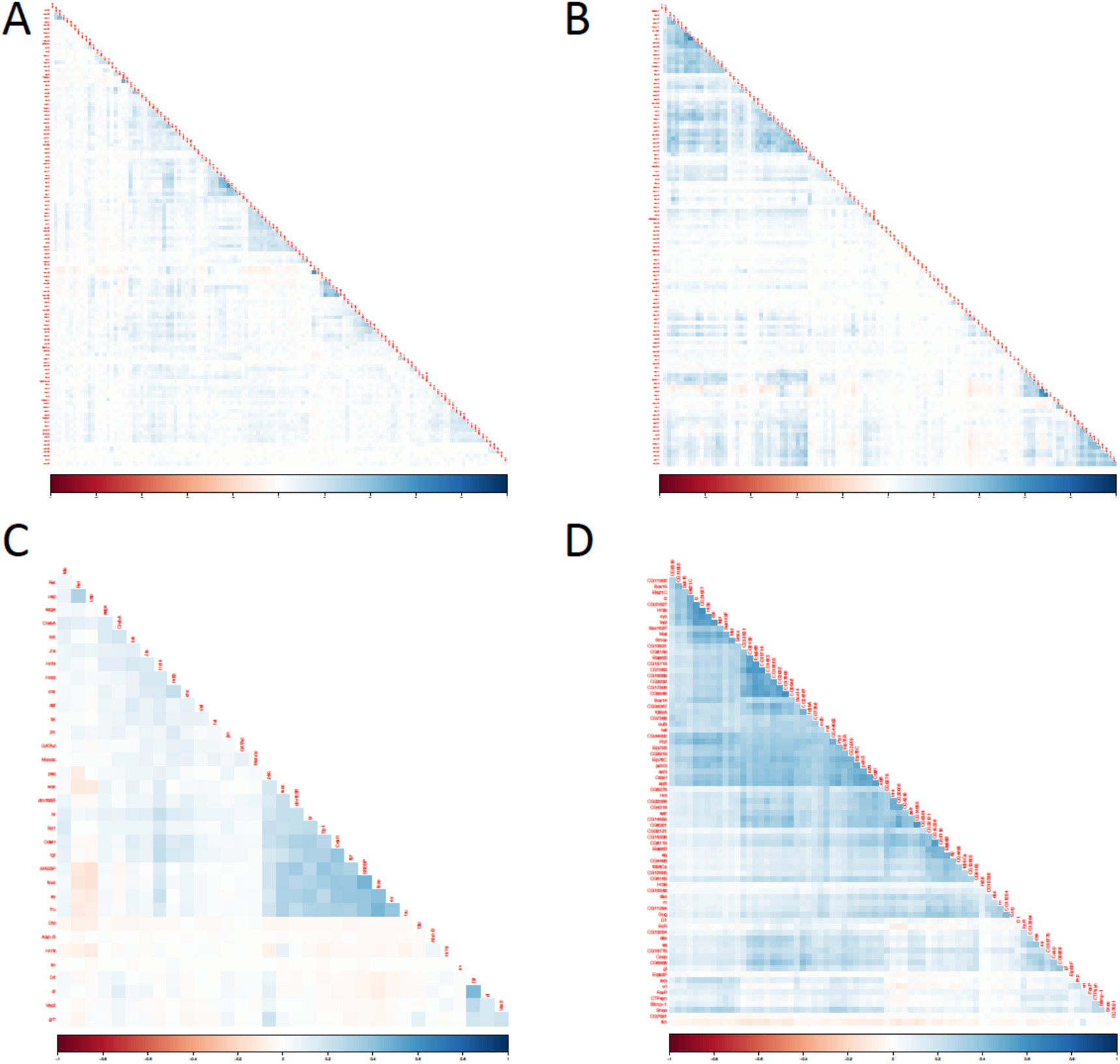
Correlation of TF-TF pairs. **A**. Pearson correlations of TFs in metapeaks in worm embryo; **B** worm larvae; **C** fly larva **D** fly pupa. In addition to the clusters of correlated TFs (blue), note also the negative correlations (red).

**Supplemental Figure 3.**
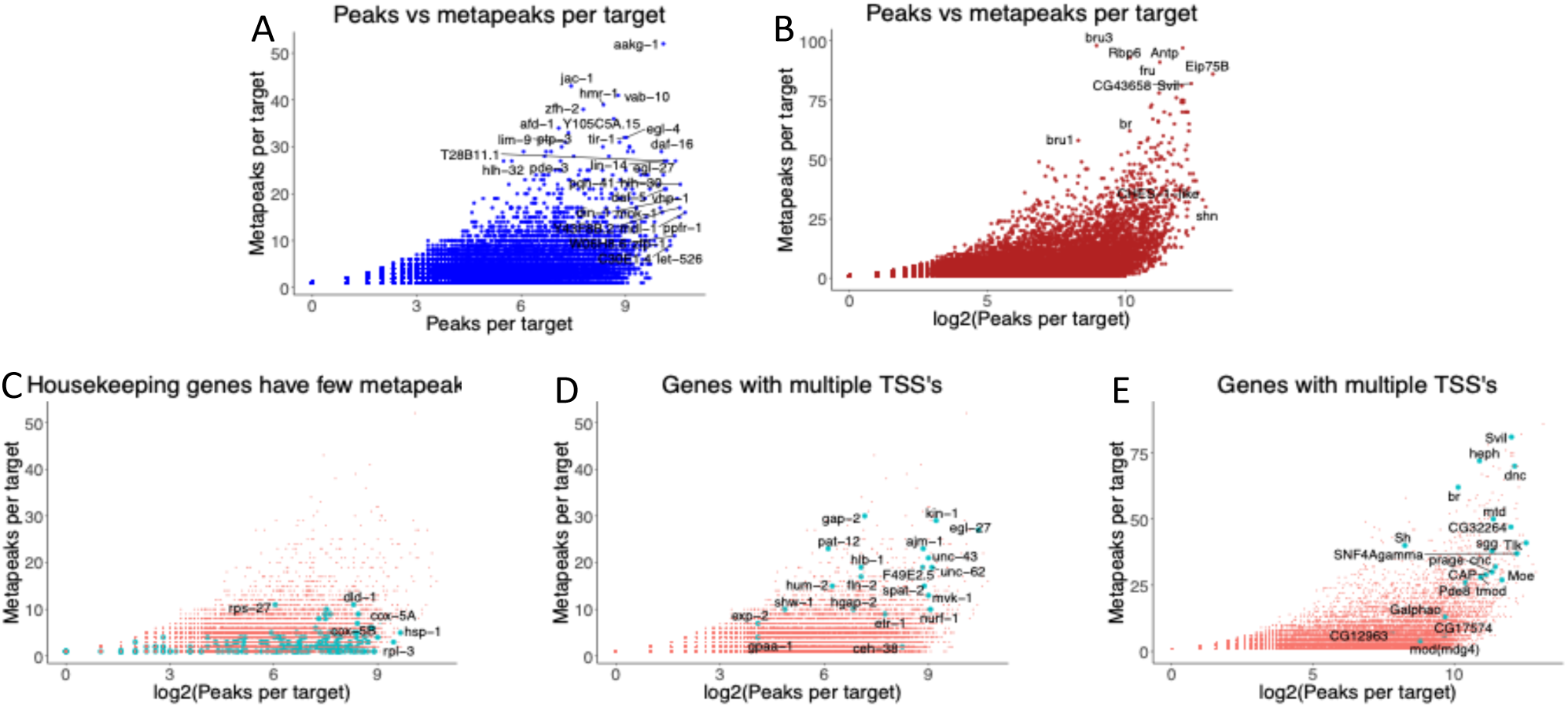
Peak-metapeak properties. **A,B** Targets in worm (A) and fly (B) can have large numbers of peaks distributed in multiple metapeaks, suggesting complex regulation. Two genes (Myo81F and Pzl) with more than 100 assigned metapeaks were omitted. **C.** In worms a small set of carefully curated housekeeping genes often have only a few, often very large metapeaks. **D,E** In both worms (D) and flies (E) genes with multiple TSS’s have multiple metapeaks, together containing high numbers of peaks.

**Supplemental Figure 4.**
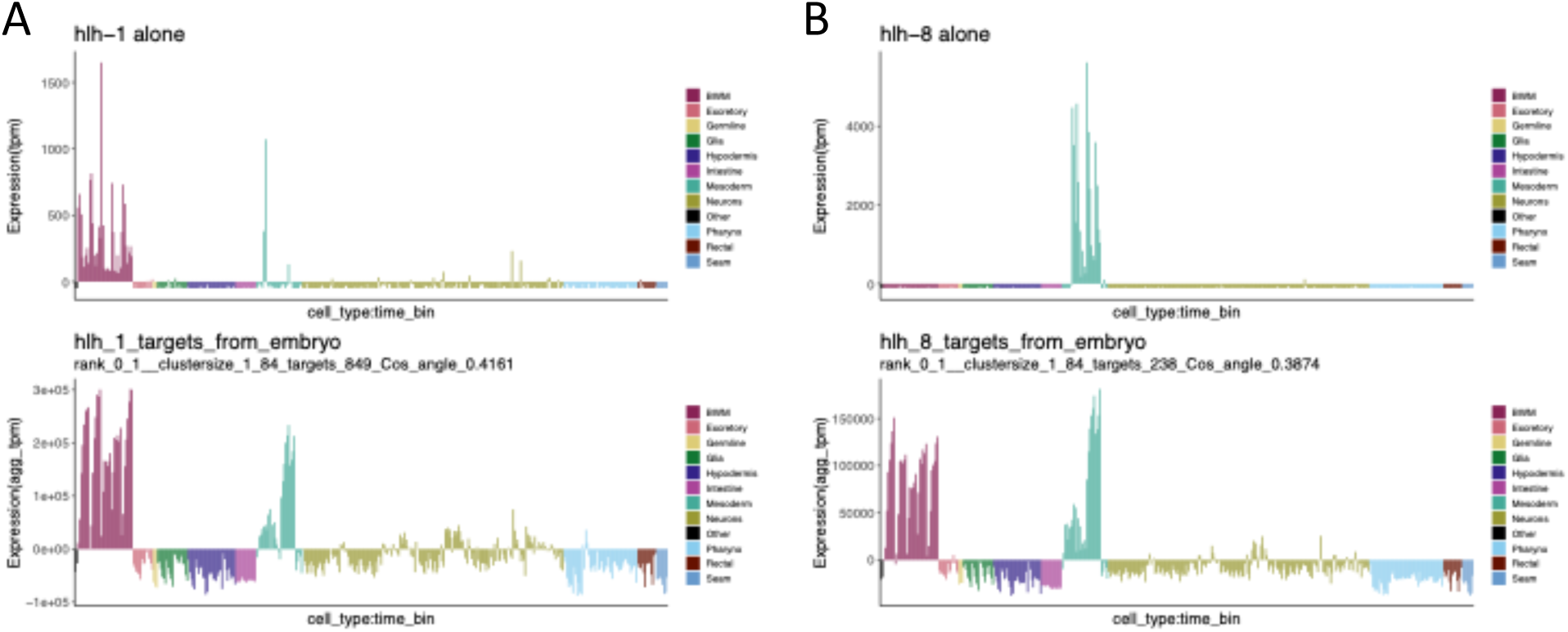
TF-target gene relationships. Some targets can be shared between different TFs. **A**The TF *hlh-1* is expressed in body wall muscles and the GLR mesodermal cells. The aggregate target expression is found there and also in other mesodermal cells. **B** The TF *hlh-8* is specifically expressed in mesodermal cells and the aggregate target expression is found there and in body wall muscle cells. The similarity of the target profiles imply shared targets and indeed *ost-1* and *pat-10*, expressed in both sets of cell types, are targets of both TFs.

**Supplemental Figure 5.**
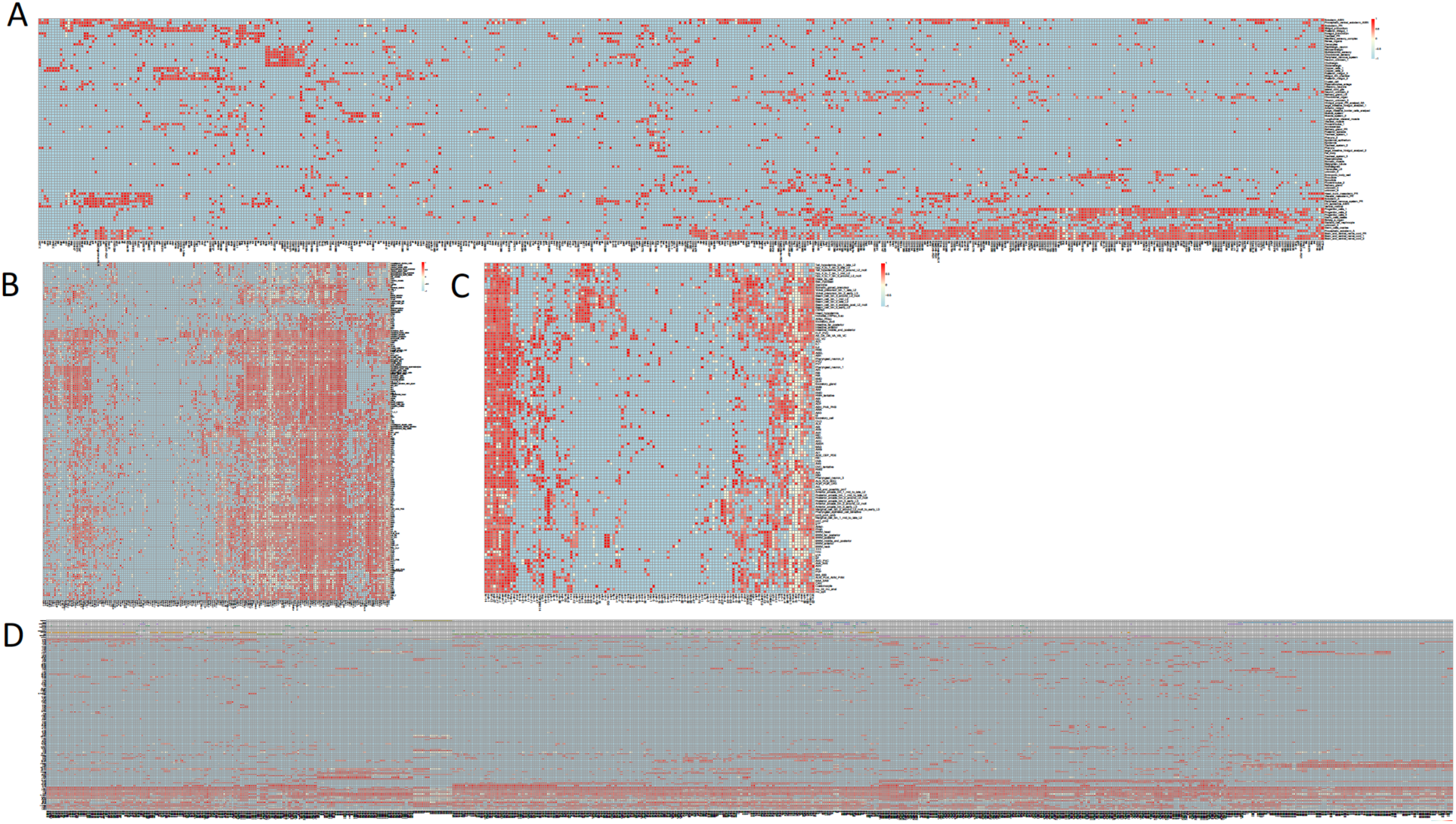
The relative importance of TFs in cell type gene expression. **A.** Heatmap showing the importance of individual TFs for predicting target gene expression patterns in single cell types for: the fly embryo; **B.** the worm larval stages; **C.** the worm L4 and adult stages; and **D.** the lineage of the worm embryo. Color scale as in Figure 7. HTML version of these maps are available on the modERN website.

**Supplemental Figure 6.**
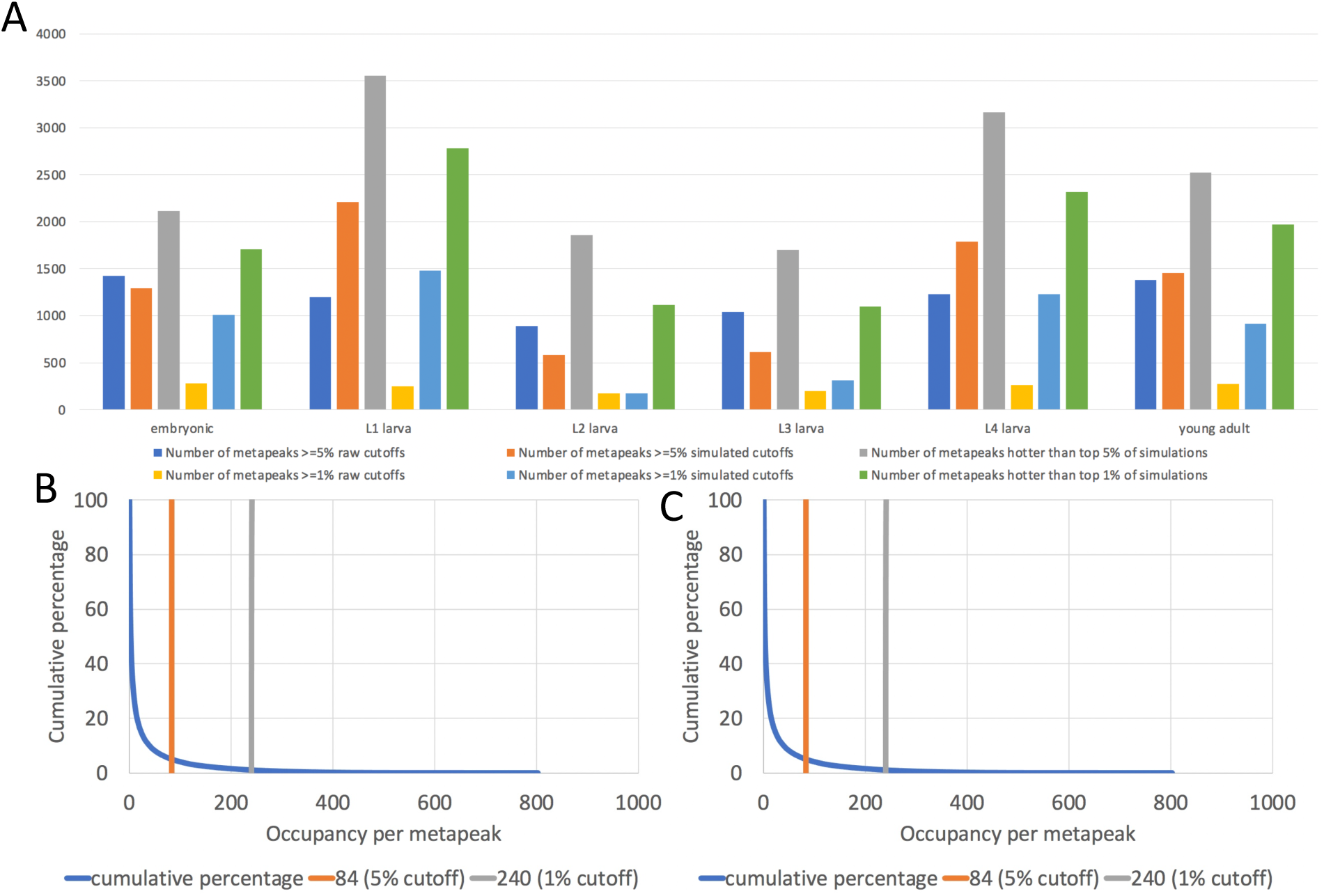
HOT site thresholding. **A.** Comparison of HOT site thresholds based on using 1% and 5% of transcription factors assayed in our data to those identified using the Araya et al (2014) approach (labeled as “number of metapeaks >=% simulated cutoffs”) and the kernel density estimation approach (L. Ma and A. Victorsen, unpublished; labeled as “number of metapeaks hotter than top % of simulations”). **B.** Decreasing cumulative percentage metapeaks as a function of occupancy per metapeak, with 5% (HOT) and 1% (UltraHOT) thresholds indicated by vertical lines (orange and gray respectively. **C**. As in B, but for fly metapeaks.

**Supplemental Figure 7.**
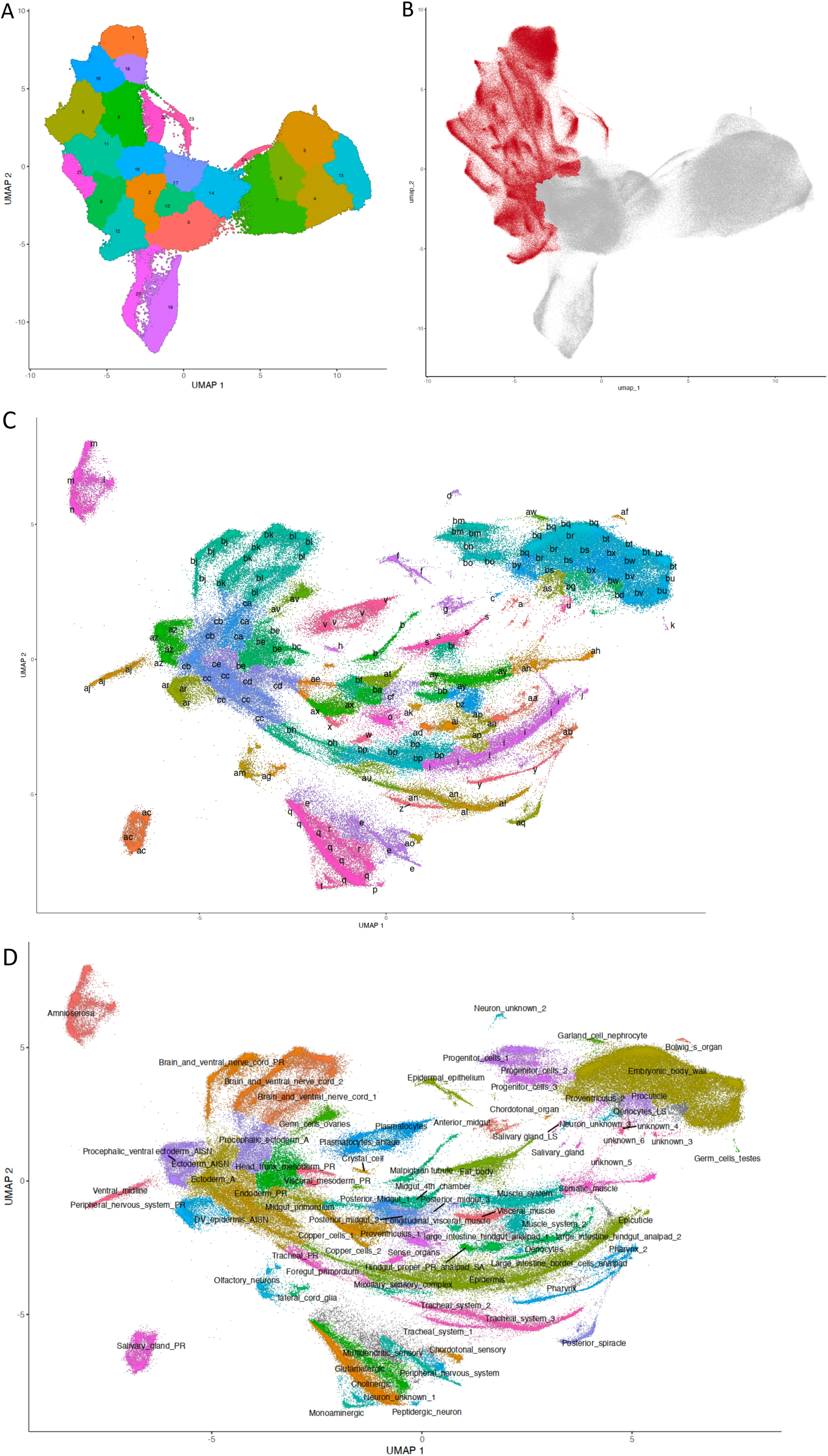
R**e**annotation **of the fly single cell RNA-seq data set. A.** UMAP of cells from Calderon et al., 2022. **B**. Older cells (red) were remapped to produce the relationships shown in C. **C**. Cells were clustered and then clusters were grouped as putative cell types and marker genes for each group were determined. **D.** The annotated cell types, determined by comparing the markers with the in situ database and literature.

**Supplemental Table 1.**
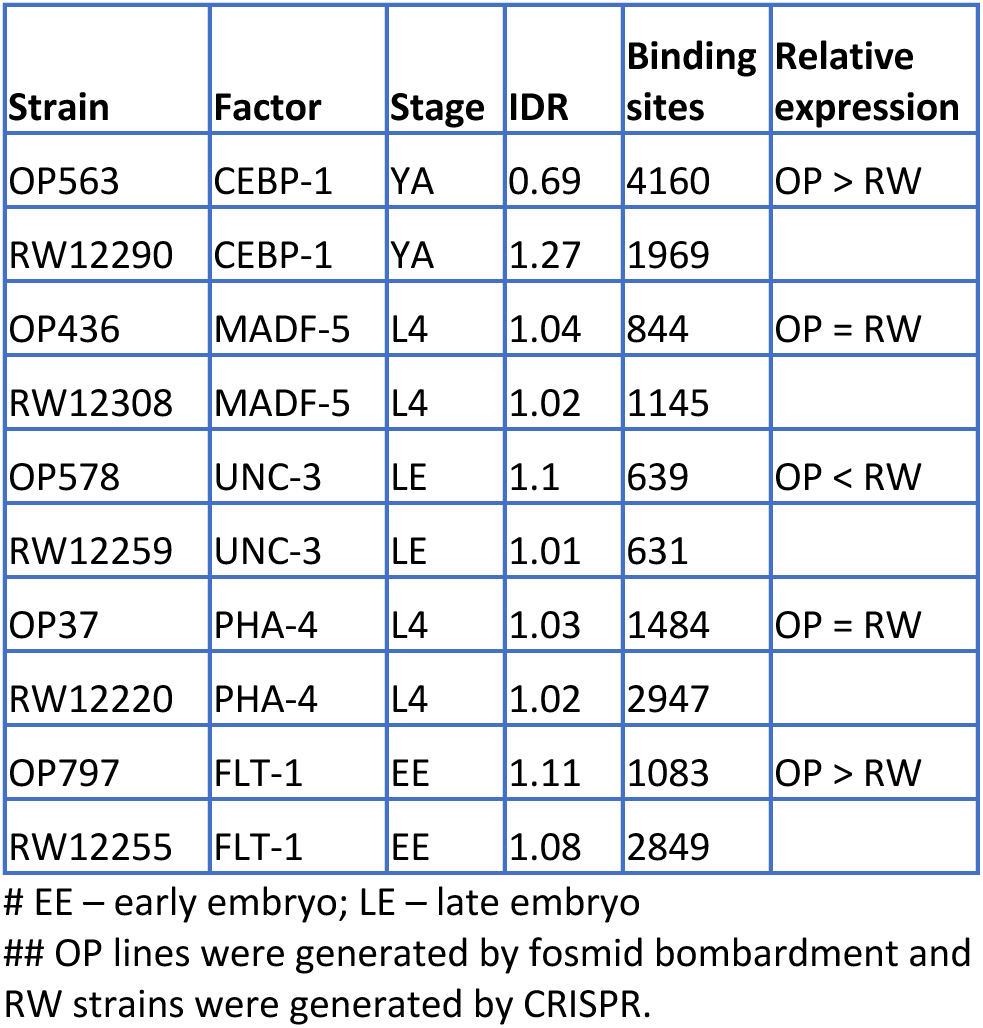

